# Doors and corners of variance partitioning in statistical ecology

**DOI:** 10.1101/2021.10.17.464682

**Authors:** Torsti Schulz, Marjo Saastamoinen, Jarno Vanhatalo

## Abstract

Variance partitioning is a common tool for statistical analysis and interpretation in both observational and experimental studies in ecology. Its popularity has led to a proliferation of methods with sometimes confusing or contradicting interpretations. Here, we present variance partitioning as a general tool in a model based Bayesian framework for summarizing and interpreting regression-like models. To demonstrate our approach we present a case study comprising of a simple occupancy model for a metapopulation of the Glanville fritillary butterfly. We pay special attention to the thorny issue of correlated covariates and random effects, and highlight uncertainty in variance partitioning. We recommend several alternative measures of variance, which jointly can be used to better interpret variance partitions. Additionally, we extend the general approach to encompass partitioning of variance within and between groups of observations, an approach very similar to analysis of variance. While noting that many troublesome issues relating to variance partitioning, such as uncertainty quantification, have been neglected in the literature, we likewise feel that the rather general applicability of the methods as an extension of statistical model-based analyses has not been fully utilized by the ecological research community either.

## 1 Introduction

In ecology, we are often interested in determining the processes and conditions that modify or maintain the distribution of species and the composition of ecological communities (Hubbell 2001, Ives *et al*. 2003, Vellend 2010, Ovaskainen *et al*. 2016). Key questions are what abiotic and biotic variables affect individual species and communities, what have been the relative roles of these variables in the past, and what will they be in the future (e.g. MacKenzie *et al*. 2007, Peres-Neto *et al*. 2006, Estay *et al*. 2014, Vuorinen *et al*. 2015, Kahilainen *et al*. 2018). These questions are intertwined but still largely separate. For example, an environmental variable such as temperature can affect species survival and reproduction rate in general but, if the study population experiences only moderate temperature variability, the contribution of temperature to variation in population size might be negligible in a particular study site (see *e*.*g*. Tomkiewicz *et al*. 1998, Veneranta *et al*. 2011). It is partially this distinction between potential variation and realized variation that has motivated our work and probably has contributed to the large array of methods of variance partitioning.

In experimental research, the relative importance of alternative variables in explaining variation in a response variable is traditionally studied using the analysis of variance (ANOVA) approach, which provides a direct method for separating background variation from treatment effects under controlled conditions (Casella and Berger 2002, Gelman 2005). In theoretical studies, the relative importance of alternative driving processes can be examined by mechanistic (first principles) models either by analytically studying their behaviour or by simulation (Ovaskainen *et al*. 2016). However, mechanistic models and large ecological experiments that would be necessary to directly study ecological processes at large scales and over longer time spans are feasible only in limited cases (*cf*. Cressie and Wikle 2011, Péron and Koons 2012, McCallum 2016). Consequently, empirical studies usually rely on observational data, often combined with model based statistical inference (Mac Nally 1996, Ives *et al*. 2003). For example, species distribution models and a variety of regression-based approaches are used to associate effects of covariates and structured random effects to the presence and abundance of species or the composition communities.

In these circumstances, where direct interpretation of effect sizes is hampered by lack of experimental control and correlations among model predictors, statistical variance partitioning methods enter the stage as an aid for interpreting the results of these regression-like models (Pedhazur 1997, pp. 241–282, Legendre and Legendre 2012). Variance partitioning is used towards various ends, such as understanding how variation is distributed at different hierarchical levels in nested experiments or multi-scale studies (Searle *et al*. 1992, Cushman and McGarigal 2002, Goldstein *et al*. 2002, McMahon and Diez 2007) and determining what is the relative importance of a covariate in terms of model performance or contribution to the process studied (Chevan and Sutherland 1991, Bring 1995, Mac Nally 1996). In other cases, the interest lies simply in separating the unique and shared contributions of covariates (Ray-Mukherjee *et al*. 2014). We also find methods such as ordination (Borcard *et al*. 1992), standardized regression coefficients (Bring 1994, 1995), variance partitioning coefficients (Goldstein *et al*. 2002), and factor analysis (Riitters *et al*. 1995) being used for variance partitioning.

The abundance of applications and methods go hand in hand with the ad-hoc nature of many approaches to variance partitioning and determination of relative importance of covariates. Often, the use of variance partitioning arises from a specific need, or for a specific kind of model. In that applied context the potential for rather wide applicability of the general approach does not become evident. These applications may also take shortcuts that do not respect the model structure or do not directly answer the question they are expected to answer (Pedhazur 1997, pp. 241–282). Furthermore, clear recommendations for how to interpret results and what should be reported when utilizing variance partitioning are lacking. To give an example, only a few studies report uncertainties in variance partition estimates or indicate the strength of correlations between variance components (but see Peres-Neto *et al*. 2006, Ray-Mukherjee *et al*. 2014, Yuan *et al*. 2017, Schulz *et al*. 2020).

As a further challenge, statistical approaches rarely fulfill the dream of causal understanding (Wright 1921, Burks 1926, Levins and Lewontin 1982, Cooper 1998, Cressie and Wikle 2011), and meaningful interpretation of estimated effect sizes of covariates and variance parameters of so-called random effects is challenging when modelling observational data. While at best the effect sizes reflect the actual importance of the covariates on the species or community studied, even then they might not reflect how important these covariates are for predicting variability in population size or community composition over space or time. The importance of the covariates in any specific context depends not only on the existence of some underlying, and usually unconfirmed, causal relationship but also on the variability in the covariates themselves.

To give a concrete example, the Baltic Sea is a brackish water basin hosting partially endemic fish species, such as Baltic herring, sprat, pikeperch, and turbot, which are all sensitive to water salinity. Even though the salinity gradient is large throughout the Baltic Sea, salinity predicts only very little of the interannual variation in the species’ reproduction rates as there is generally little interannual variation in water salinity in regions where these fish species occur (e.g. Lappalainen and Lehtonen 1995, Casini *et al*. 2006, Nissling *et al*. 2006). It is also often the case that any of the survey data at hand do not represent the full extent of variation in the covariates in the areas where the species occur (Vernier *et al*. 2008, *cf*. Foster *et al*. 2021). In either case, if data contain little or no variation with respect to a covariate, then there is no statistical way to assess its effect and thus no variation in the outcome can be attributed to that covariate, whether it is important or not (Pratt 1987). In extreme cases the situation is akin to the lack of genetic variability in an allele fixed either due to its high fitness advantage or simply due to genetic drift (*cf*. Whitlock and Gomulkiewicz 2005).

Keeping the preceding admonitions in mind, we follow Gelman (2005, Gelman *et al*. 2014, 381– 400, Gelman *et al*. 2019) to present a model based approach to variance partitioning. Our goal is to quantify the contributions of individual model components to variation in the quantity of interest, such as species abundance or community composition. A similar Bayesian approach has been utilized by *e*.*g*. Yuan et al. (2017), who used it as a model checking tool for a spatio-temporal species distribution model, by Schulz *et al*. (2020) where we used it to explore occupancy–abundance relationships in a butterfly metapopulation, and in the joint species distribution modelling software HMSC (Ovaskainen and Abrego 2020, Tikhonov *et al*. 2020). Instead of focusing only on a specific application, we present variance partitioning as a more general tool for the analysis and interpretation of statistical models. We also demonstrate how we can avoid some of the pitfalls of conducting and interpreting variance partitioning studies present in earlier work as well as show its applicability to predictive scenarios. We specifically address topics such as correlations between covariates, confounding by random effects and correlated parameters, which have received only little attention in the contemporary literature (but see Wakefield 2003, 2004, Yuan *et al*. 2017). Moreover, we propose variance partitioning for grouping of covariates as well as conditioning on observations and present some common as well as new visualization techniques for variance partitions. We derive and discuss the methods in a generalized linear model setting but variance partitioning is, at least partially, applicable to a wider class of statistical models, namely any hierarchical additive models (Gelman 2005), including traditional GLMs and GAMs as well as their extensions with structured and unstructured random effects (Hodges *et al*. 2007).

The rest of the paper is organized as follows. We first introduce the basic concepts behind variance partitioning in Section 2.1, after which we introduce the theory for its extensions in sections 2.2–2.7. In Section 3, we demonstrate the different variance partitioning techniques with a case study using a simple species distribution modeling (SDM) -like regression model. We will apply it to model presence–absence of an insect herbivore in a metapopulation context. After Section 2.1, practically oriented readers can jump directly to the case study and refer to the theory after reading the motivating examples. For researchers interested in applying these methods we also provide worked examples including all necessary code in the Programming supplement.

## 2 Material and methods

### 2.1 Introduction to variance partitioning

To better understand how variance partitioning works and is easiest to interpret, we first look at an idealized scenario from population ecology, where a closed population reproduces according to a simple time-varying Ricker population model (Ricker 1954). The Ricker model for population growth at time *t* with population size *N*_*t*_ can be written as (Ruppert and Carroll 1985, Collie *et al*. 2012, Weigel *et al*. 2021)

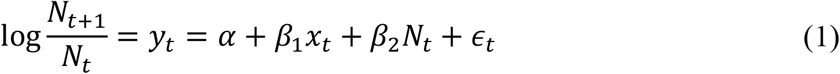

where *α + β*_1_*x*_*t*_ stands for the intrinsic population growth as a function of environmental covariates *x*_*t*_, *β*_2_ (< 0) is a parameter governing density-dependent survival and *∈*_*t*_ models intrinsic stochasticity in the population dynamics. Thus, we can describe the log growth rate *y*_*t*_ of this idealized population with a linear model, whose parameters have an intuitive ecological interpretation. Hence, we would hope calculating and interpreting the variance partitioning were straightforward as well. However, this is not the case in practice.

Let us first assume that we know the parameters of the model, *θ* = [*α, β*_1_, *β*_2_], and we can measure the population size and environmental covariates accurately for years *t* = 1, …, *n*. The standard approach to analyze what proportion of the variation in log-population growth rate arises from the three different processes described in our model would be to calculate

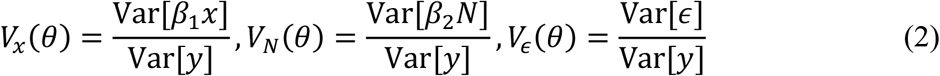

where the variances are sample variances over the *n* observations collected in vectors, such as *y* = [*y*_1_, …, *y*_*n*_]^T^.

In order for these terms to summarize the *proportions* of variation they should sum up to one. However they sum up to one, only in the special case that the three components of the Ricker model are mutually independent so that Var[*y*] = Var[*β*_1_*x*] + Var[*β*_2_*N*] + Var[*ϵ*]. This is an unrealistic assumption in most cases since, for example, temporal correlation in environmental covariates (Cov[*x*_*t*_, *x*_*t*−1_] ≠ 0) induces correlation between the intrinsic growth rate and the density dependent population dynamics (Mathematical supplement 1, see also Ives *et al*. 2003). If the correlation between *β*_1_*x*_*t*_ and *N*_*t*_ is positive then Var[*β*_1_*x*] + Var[*β*_2_*N*] + Var[*ϵ*] < Var[*y*] and if they are negatively correlated, then the sum of variance terms is larger than the variance of *y*.

In ecological applications, it is common to have temporally or spatially correlated environmental covariates and we refer to this phenomenon as the *correlated covariates challenge* (see also Cox and Snell 1974, Pedhazur 1997, pp. 241–282, Graham 2003, Dormann *et al*. 2013, Morrisey and Ruxton 2018). The example of fisheries in the Baltic Sea above serves to illustrate the problem: salinity, water temperature, and depth can all play a role in determining spawning areas for many of the fish species, but they are hard to disentangle as they are also all highly correlated with each other (Tomkiewicz *et al*. 1998, Janssen *et al*. 1999, MacKenzie *et al*. 2007, Veneranta *et al*. 2016).

A related challenge that has received less attention in the ecological literature is the challenge posed by correlation between the random effects and the covariates (Wakefield 2004, Yuan *et al*. 2017). The random effects *∈*_*t*_ are typically defined to be *a priori* independent from the other model components. However, most often our model does not contain all relevant covariates, but we may have missed a covariate *z*_*t*_ that is correlated with *x*_*t*_. When estimating the random effects from data, the effect of this missing covariate will be captured by *∈*_*t*_ at least partially, leading to correlation between *β*_1_*x*_*t*_ and *∈*_*t*_. Correlation between *β*_1_*x*_*t*_ and *∈*_*t*_ can also arise for other reasons such as when a model includes both a spatial random effect and covariates with spatial structure (Teng *et al*. 2018). We call this phenomenon the *confounded random effects challenge*.

One approach to surpass the correlated covariates and confounded random effects challenges is to calculate the variance partitioning as

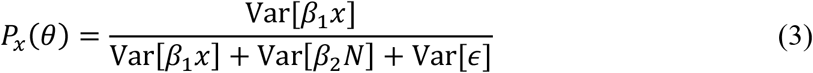

which ensures that the different variance components sum up to one by ignoring the covariances in the denominator (Ovaskainen and Abrego 2020, pp. 69–70). We call this the *diagonal variance partition* as when looking at the covariance matrix of the components, this measure only accounts for the diagonal variation. The diagonal variance partition is only valid when the correlations are negligible or of no consequence to the interpretation of the results, so it is at most a good approximation for a ‘true’ variance contribution of the covariate. Hence, this diagonal approximation is not a general solution to the problem.

Up to this point, we have assumed that the model parameters are known. Naturally, this is usually not the case, since if it was, we would not need to do statistical inference in the first place. In this work, we follow the Bayesian approach and calculate the full posterior distribution for model parameters *p*(*θ*|*D*) where 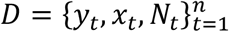 collects the data and *θ* = [*α, β*_1_, *β*_2_] the parameters, as above. The posterior distribution for the parameters induces a posterior distribution for the variance partitioning statistics, so that instead of a single estimate for *V*(*θ*), the variance partitioning, we have a random variable (or rather random vector) with distribution *V ∼ p*(*V*|*D*) (*cf*. Gelman *et al*. 2019). From a Bayesian point of view, it is natural to study this posterior distribution which we can use as a measure of the uncertainty in our variance partitions. The uncertainty in variance partitioning is not, however, typically reported but what is traditionally done (e.g. Cushman and McGarigal 2002, Legendre and Legendre 2012) is that people summarize the model parameters with a point estimate 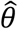, such as the posterior mean, or more commonly the least squares or maximum likelihood estimate, and report 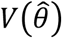 or 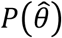 or some similar measure (but see *e*.*g*. Goldstein *et al*. 2002, Yuan *et al*. 2017). These estimates suffer from the correlated covariates and confounded random effects challenges as discussed above.

However, there is even more to the story. In addition to neglecting the uncertainty in variance partitioning, there is no guarantee that 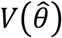 and 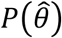 are actually good summaries for *p*(*V*|*D*) and *p*(*P*|*D*). Even if the parameters were uncorrelated *a posteriori*, directly substituting point estimates into the formulas will be erroneous. Take for example the variance partition of the covariate effect *V*_*x*_(*θ*) ∝ Var[*β*_1_*x*] (the ∝ denotes “is proportional to”). The plug-in estimate 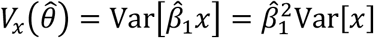 where 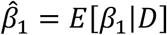 (the posterior mean) would not lead to the posterior mean E[*V*_*x*_|*D*] since 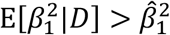 unless the variance of *β*_1_ is zero (Jensen’s inequality). In the case of *P*_*x*_ the difference between the plug-in estimate 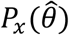 and E[*P*_*x*_|*D*] is typically even larger, since also its denominator is a function of parameters (see eq. (3)). This point is especially salient in the Bayesian context, but can apply equally to some frequentist measures of uncertainty about the point estimate of variance partitioning.

The correct approach to calculate a point summary for the mean and spread of a variance partition is using the distributions *p*(*V*|*D*) and *p*(*P*|*D*), most generally using the expectation and variance of their distribution

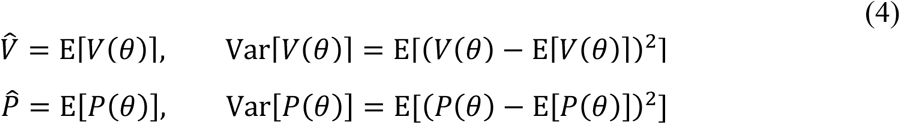

These summaries can easily be approximated using samples from the posterior distribution of *p*(*θ*|*D*) as it is sufficient to know the distribution of the parameters *θ* to calculate the expectation of functions of these parameters (Casella and Berger 2002, p. 55; see also Section 3, Mathematical supplement 2.7, and the Programming supplement). This approach avoids the bias that substituting point estimates into the formulas directly introduces and is used for example in HMSC (Table 1). However, if we then only communicate the posterior means 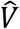 or 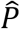, we lose knowledge of how the uncertainty in model parameters propagates to the variance partitions. The least should be to also report the posterior variance of the estimates.

**Table 1.** Notation and terminology for finite population variance partitioning measures.

| Notation | Concept | Comment |
| --- | --- | --- |
| $V_x$ | *Variance partition* as a random variable | Proportion of variation in the linear predictor due to the linear term $x$. |
| $P_x$ | *Diagonal variance partition* as a random variable | Proportion of independent variation in the linear term $x$. |
| $V_x(\hat{\theta})$ | Variance partition at a point estimate | Biased estimate of $V_x$ calculated from a point estimate of the parameters. |
| $P_x(\hat{\theta})$ | Diagonal variance partition at a point estimate | Biased estimate of $P_x$ calculated from a point estimate of the parameters. |
| $\hat{V}_x$ | Point estimate of the variance partition | Location point estimate (e.g. posterior mean) of the distribution of $V_x$. |
| $\hat{P}_x$ | Point estimate of the diagonal variance partition | Location point estimate (e.g. posterior mean) of the distribution of $P_x$. |
| $\text{Var}[V_x]$ | Posterior variance of the variance partition | Summary of the posterior uncertainty in the variance partition estimate. |
| $\text{Var}[P_x]$ | Posterior variance of the diagonal variance partition | Summary of the posterior uncertainty in the diagonal variance partition estimate. |
| $b_l$ | *predictor group* effect in generalized linear regression | Linear combination of the linear terms belonging to the group $l$. |
| $V_x$ | *Sample variance partition* | Variance partition calculated over the observations included in the study |
| $V_x^{\text{pop}}$ | *Population variance partition* | Variance partition calculated over the population of which the observations are a sample. |
| $V_x^{\text{pred}}$ | *Predictive population variance partition* | Variance partition calculated over some new, possibly hypothetical, set of observations. |

In the Bayesian context, we must also consider what happens when the model parameters are mutually correlated in their posterior distribution, that is, for example, Cor[*β*_1_, *β*_2_|*D*] ≠ 0. The posterior correlations between parameters translate into posterior correlations between the variance partition measures *V*_*x*_ and *V*_*N*_ or equivalently *P*_*x*_ and *P*_*N*_. In that case, even the marginal distributions of the variance partitions are not necessarily good summaries of their joint posterior distribution (see Sections 2.6 and 3.6).

### 2.2 Variance partitioning of the linear predictor

Moving on to a general regression context, we discuss variance partitioning in the context of the *linear predictor*. In a generalized linear model, the linear predictor is *η* = *g*(*E*(*y*)), a transformation of the expected value of outcomes *y* to the linear space using a link function *g*. Linear predictor is a length *n* vector, that is formed as the sum of *d* vectors of *linear terms*, that is 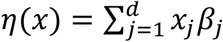 where, 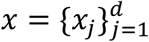 and *x*_*j*_ is a length *n* vector of covariates. But as far as variance partitioning is concerned the linear terms could equally be additive or random effects. The variance over the *n* observations of the linear predictor in (generalized) linear models is 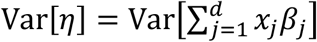. Hence its decomposition follows from the properties of variances of linear combinations (Mulaik 2010, pp. 73–76, 83–84) and can be written as a sum of the covariances of the linear terms:

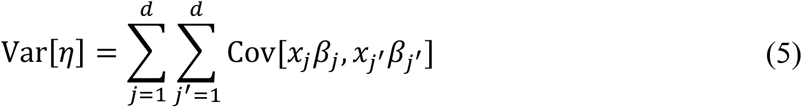

Here our variance partition measures are 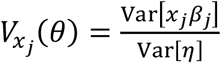 and 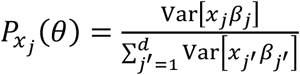 for the *j*th covariate *x*_*j*_. To see how this generalizes to any generalized additive model with random effects, remember that if we write the linear terms in the form *f*_*j*_(*x*_*j*_) then 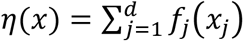, and we have an unweighted linear combination of the additive functions or random effects and hence can still apply all the methods presented in this paper (i.e. 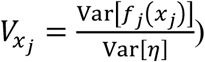). We could also calculate the variance partitioning for non-linear joint effects of two or several covariates, such as *f*(*x*_*j*_, *x*_*j*′_). This is used, for example, when calculating the variance contribution of continuous spatial random effects.

The approach is not limited to partitioning the variation in the linear predictor, but can also be used for the more traditional partitioning of explained variation, *R*^2^, when working with a linear model or a model with linearizable residuals. If we partition variance while accounting for residual variance, it will give us a measure that is equal to the traditional *R*^2^ when defined as 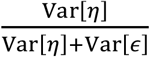. The only change for the variance partitioning metrics *Vx* (*θ*) and *Px* (*θ*) is the inclusion of the residual variance in the denominator so that 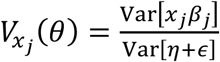 and 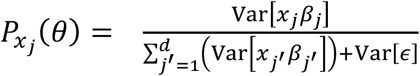. The residuals *∈* can be calculated either with respect to the observations *y* or its posterior predictive distribution 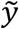 as is done in the Bayesian *R*^2^ approach (Gelman *et al*. 2019) but we do not consider this choice further here.

### 2.3 Grouping linear terms in variance partitioning

Often analyses include many environmental covariates, which can be divided into multiple groups, such as habitat quality or climatic variables. These *groups of linear terms* are sometimes called *predictor groups* (Ovaskainen and Abrego 2020, pp. 69–70) or simply *sets of variables* in some ecological literature (e.g. Legendre and Legendre 2012, *cf*. Wisler 1969, Beaton 1973, Cox and Snell 1974). We can calculate the variance partitioning at the level of these groups of linear terms by summing over the variance terms of the covariates included in that group (*cf*. Mulaik 2010, pp. 72–84). To present this explicitly, we define the following notation. If each covariate is associated with only one of *m* groups, the linear term group effects can be written formally as 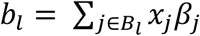 where *l* = 1, …, *m* and *B*_*l*_ is a set of covariate indexes that are associated with the *l*’th group. We can rewrite the linear predictor in terms of sums of the linear term group effects, 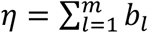. The variance of a group effect is 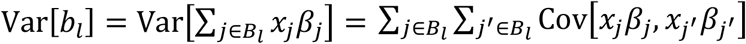 and can be used to express the variance of the linear predictor 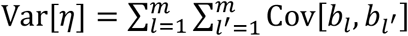. The linear predictor group variance partition is then

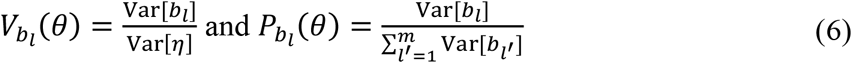

for some group *b*_*l*_ (Mathematical supplement 2.3).

In addition to simplifying the presentation of models with many covariates by summarizing them at a more abstract thematic level, it can also help interpreting models with correlated covariates when the correlations between covariates are greater within the linear term groups than between them: grouping related covariates incorporates the covariances between those covariates to the variance term of that group and thus reduces the covariance between the (grouped) variance components (Ovaskainen and Abrego 2020, pp. 69–70). This can also reduce the posterior correlation between the variance partitions (see Section 2.6) compared to a situation without groups. However, this does not solve either of the problems entirely, since correlations can occur also between thematically unrelated covariates leading to covariances and posterior correlations between the groups.

It is to be noted, that the grouping of the covariates is not a property of the observations, or the current sample of data at hand, but rather a choice made by the analyst as will be illustrated in the case study (Section 3.1). It thus incorporates external knowledge into the interpretation of results. Also, depending on how the groups of linear terms are defined, a group might consist of a mixture of covariate and random effects (*cf*. Schulz et al. 2020).

### 2.4 Variance partitioning over populations and its use in prediction

The approach to variance partitioning presented thus far corresponds to partitioning with respect to the observations at hand, such as data from specific time points and sites. Hereafter, we refer to them as *sample variance partitions*, since they describe the properties of a finite sample from some larger domain, for example a set of potentially randomized sampling locations in a larger landscape or sampling times of some continuous process. In some cases, it will be possible to partition variance with respect to the full domain of interest (see Section 3.3). The domain can, for example, be the area and time period over which a survey was conducted (see Yuan *et al*. M2017). This *population variance partition* will often correspond closer to what is of ecological interest, especially when modelling observational data obtained without a well-designed survey. In such data, the distribution of environmental covariates in the observations does not correspond to that of the whole study region and the *sample variance partition* will not be representative. However, whether population variance partitions can be calculated or not, depends on the availability of covariate data over the domain and is thus often not easily realizable. The distinction between sample and population variance partitioning is demonstrated in the case study (section 3.3) and Fig. 2 ((a) and (b)).

The notion of variance partitioning over an existing, but only partially observed region or population, can be extended to *predictive variance partitioning* over new domains. For example, in climate change studies, we could be interested in predicting how much variation in species distribution is contributed by the environment under future climate compared to the current climate. When predicting variance contributions for a new location or time, the distribution, and hence variability, of the predictors often differs from the original study location or time, even if in terms of the range of values for individual covariates the prediction would have the character of “interpolation”. Hence, population variance partitioning can also be a tool for summarizing ecological predictions and scenarios.

### 2.5 Conditional variance partitioning and partitioning between and within conditions

If we group our observations according to some categorical variable or some function that assigns observations to groups based on the covariates, we can calculate variance conditional on that grouping. For example, our case study (Section 3.4) includes a covariate for the abundance of the insect herbivore’s host plant. It is divided into three categories, with all observations within the same abundance category grouped together. Formally, we define a conditioning function *g*(*x*) that assigns each observation into one of *G* predefined groups. The variance partitioning conditional on group *c ∈ G* is then calculated over the subset of observations (or the subset of the population) for which *g*(*x*) = *c*. For example, the conditional sample variance partitioning for the covariate *x*_*j*_ would be

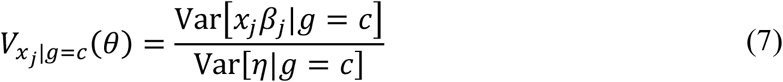

where *Var*[· |*g* = *c*] denotes the sample variance over the subset of observations for which *g*(*x*_*i*_) = *c* (*i* = 1, …, *n* indexes the observations, see also Mathematical supplement 2.4). The definition for 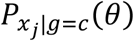 would arise analogously.

When, for example, a predictor varies only over some subset of the observations, conditional variance partitioning can be used to study its role in that subset where it varies. However, conditional partitioning should be used with some meaningful question in mind, not just to search for an arbitrary division of the data to make up for the minor role of a predictor for the whole data. Unlike the groups of linear terms, the conditioning variable is a property of the data – that is the group membership of each observation depends on the values of the covariates (or functions thereof) available for the observations. Usually these variables would be part of the covariates of the linear predictor. Nonetheless, the choice of the conditioning variable is still made by the analyst.

We can extend conditional variance partitioning to see how much of the total variation is partitioned within the groups defined by conditioning variable and how much variation is due to the differences between the conditions. This approach has the character of analysis of variance (*cf*. Pedhazur 1997, pp. 679–683, Nakagawa and Schielzeth 2013). It utilizes the law of total (co)variance Var[*x*_*j*_*β*_*j*_] = E [Var[*x*_*j*_*β*_*j*_|*g*]] + Var [E[*x*_*j*_*β*_*j*_|*g*]] to split total variation into variation within (E [Var[*x*_*j*_*β*_*j*_|*g*]]) and between (Var [E[*x*_*j*_*β*_*j*_|*g*]]) conditions (Whittaker 1990, pp. 124– 125).

Specifically, in applied variance partitioning we utilize the sample covariance of the linear terms within and between conditions. The variance partition within conditions is simply the weighted average of the conditional variances

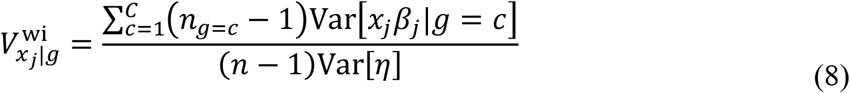

where *n*_*g*=*c*_ is the number of observations belonging to the condition *c* and 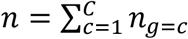 is the total number of observations. The variation between groups is derived from the weighted variance of the per-condition means of the linear terms

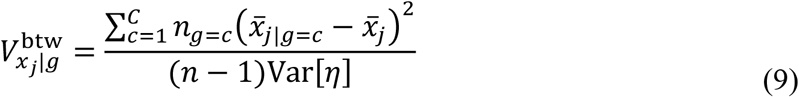

where the 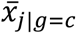 is the per-condition mean and 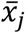 is the unconditional or grand mean of the linear term over all *n* observations. The within and between condition diagonal variance partitions 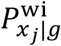 and 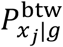 are defined analogously. Note that the sum of the variance partitions within and between conditions equals the variance partitioning over all observations, i.e. 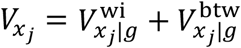. For derivation of the within and between condition variances see the Mathematical supplement 2.5.

Differences between the conditional variance partitions imply that the distributions of covariates and random effects differ between the conditions. The partitioning of variation to within and between condition (co)variance is a way to summarize these differences for individual linear terms or predictor groups. When the conditioning variable is a categorical predictor or the level of an *iid* random effect, then the corresponding linear term will contribute only to between-condition variation. At the other extreme, a predictor that is constrained to sum to zero within each condition would only contribute to within-condition variation.

### 2.6 Correlation between covariates and posterior correlation between parameters

The variance partitions *p*(*V*|*D*) and *p*(*P*|*D*) typically include correlations between their components, the *V*_*x*_s and *P*_*x*_s (see also Fig. 3) – an issue which is often overlooked when dealing only with point estimates. These correlations are a function of both correlations among the covariates and the posterior correlations among the model parameters *θ*. Generally, whenever one variance component has a higher contribution, it must be offset by a reduction in the other components. How this reduction is distributed among the other components depends on the correlations between the covariates and model parameters. The correlation between variance components contributes to uncertainty in variance partitioning; that is, the role of any single component cannot be independently assessed in the presence of such correlations, just as with the covariate effects themselves. Only when the correlations are low is it always safe to work directly with the marginal distributions of the individual variance partitions. As noted above, grouping of linear terms often ameliorates this issue, as the within-group correlations will be subsumed by the marginal distribution of the grouped variance partitions (*V*_*b*_; Section 2.3). Since this does not totally remove the problem, it is recommendable to explicitly report and discuss the correlations between different variance components (see Section 3.6).

The problem caused by the correlations is analogous to an over a century old problem of how to interpret variance partitions in a regression context in the presence of correlations between the model’s covariates (Wright 1921, Burks 1926, Cox and Snell 1974, Bring 1995, Graham 2003, Dormann *et al*. 2013). Hence, by exploiting this literature, in addition to the measures *V*_*x*_ and *P*_*x*_ presented above, we present two other measures which can be used to summarize the covariance relationships between the linear terms and can also be interpreted as variance partitions themselves (Table 2). To simplify the notation we denote here the linear terms by *a*_*j*_ = *x*_*j*_ *β*_*j*_ such that 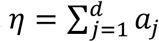. We begin by rewriting the variance of the linear predictor in two alternative ways

**Table 2.** Various variance partitioning measures derived from the covariance matrix of the linear terms. The *Sum* column indicates the sum of the measure over all linear terms. The *Value* column shows the range of values the measure can take for a single linear term; an empty cell means there is no constraint on the value. An “x” in the *Covariance* column indicates that the measure is a function of the covariance between the linear terms. Note that in the context of model inference, even *P*_*x*_ is of course affected by the covariance but the variance measure itself can be calculated without referencing the covariance. In the case of uncorrelated linear terms all measures would equal each other.

| Measure | | Sum ($\sum_{j=1}^d \cdot$) | Value (·) | Covariance | Description |
| --- | --- | --- | --- | --- | --- |
| Variance partition | $V_x$ | $\geq 0$ | $\geq 0$ | x | Proportion of variation due to the linear term |
| Diagonal variance partition | $P_x$ | 1 | $[0, 1]$ | | Proportion of diagonal variance due to the linear term |
| Unique variance partition | $U_x$ | $[0, 1]$ | $[0, 1]$ | x | Proportion of variation uniquely due to the linear term; square of the semi-partial correlation coefficient |
| Marginal variance partition | $M_x$ | 1 | $[-d + 2; d]$ | x | Marginal or averaged variance contribution of the linear term; equals the covariance between the linear term and the linear predictor. |

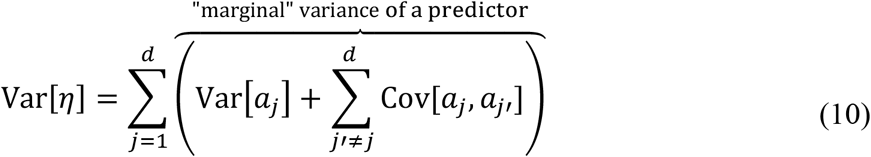

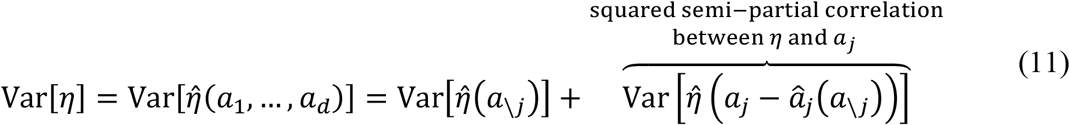

where *a*_\*j*_ collects all other linear terms except *a*_*j*_ and the hat-notation denotes linear least squares predictor so that, for example, 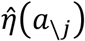 is the linear least squares prediction for *η* using *a*_\*j*_; since 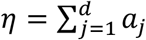, it holds that 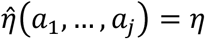. The first equality above arises by just rearranging the covariance terms in (5) and the second equality is proved for example by Whittaker (1990, pp. 134–138). In linear regression, the sum of the terms inside parenthesis on the right hand side of (10) we call the marginal variance of the corresponding linear term since it is a measure of the contribution of a linear term, while accounting for covariances with all the other linear terms (Pratt 1987, Chase 1960, Bring 1995). The squared semi-partial correlation coefficient between the linear term, *a*_*j*_, and the linear predictor, *η*, in (11) measures the variance contribution of the linear term, that cannot be accounted for by a linear combination of the other linear terms (Whittaker 1990, pp. 134–137, Legendre and Legendre 2012, pp. 173–180). Hence, squared semi-partial correlation coefficient between a linear term and a linear predictor can be seen as the unique variance from that linear term.

Motivated by the above considerations we define the *marginal variance partition* as

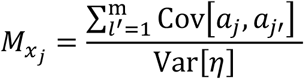

and the *unique variance partition* as

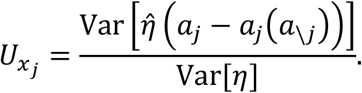

The marginal and unique variance partitions, *M*_*b*_ and *U*_*b*_, for a group of linear terms are calculated analogously. The posterior distribution for both of these measures is remarkably simple to calculate by exploiting the properties of the sample covariances between the linear predictor and individual predictive terms and is easy to extend to groups of linear terms as will be illustrated in Section 3.6 and described in detail in the Mathematical supplement(2.6).

The unique variance partition is equal to *V*_*x*_ only in the absence of correlations between the linear terms. In all other cases, its value is less than the value of *V*_*x*_ and it is zero if the linear term under investigation can perfectly be explained by the other linear terms. Hence, the difference between *U*_*x*_ and *V*_*x*_ is a measure of the effect of correlations between linear terms to the variance partition (see also Burks 1926, Whittaker 1990, pp. 137–140). *U*_*x*_ on its own can be thought of as a lower bound of variability associated with that linear term. In the case of treatment effects in an experimental context, it would be the part of variation in the outcome that is unequivocally attributable to the treatment corresponding to that linear term (i.e., the direct effect). In the case of auxiliary variables or observational data, it simply implies that that proportion of variation in the linear term is not confounded with the other linear terms in the model. However that “unique variation” could still be confounded with covariates missing from the model. We note also that, even though in some special cases the semi-partial correlation can be interpreted as a causal effect (*cf*. Burks 1926), that is not generally the case and, hence, we should not, in general, give causal interpretation for the unique variance partition.

Unlike the unique variance partition, the marginal variance partition can give information about the effect and sign of covariances with other linear terms. The unscaled measure, 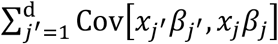, is also equal to the covariance between the linear term and the linear predictor (see Mathematical supplement 2.6.3). Hence, when the measure is close to zero, that linear term has no direct effect on the linear predictor, only an indirect effect via its correlations with the other terms. That indirect effect is sometimes interpreted as improving the model due to removal of “noise” from other covariates (Bring 1995). At the same time, the marginal variance partition can also equal the variance partition *V*_*x*_ even in the presence of covariances, if they “cancel out”. The marginal variance partition shares some properties with commonality analysis, a widely used method of variance partitioning in ecology (Legendre and Legendre 2012, 570– 583), namely it sums up to one, represents the direct effect of the linear term and comprises the unique and shared variance contributions of that linear term (Chase 1960, see also Beaton 1973, Mathematical supplement 2.6.4).

We illustrate possible conclusions from comparing different values of these three measures for a single linear term or a group of linear terms in Table 3 and present some of them here: When the marginal variance partition *M*_*x*_ is larger than the variance partition *V*_*x*_, this implies that the covariances between linear term *x* and the other linear terms are positive on average and increase the variability of the linear predictor *η*. When *M*_*x*_ is smaller than *V*_*x*_, the covariances are negative on average. If *M*_*x*_ is still positive there is some confounding between the linear terms, but the linear term still contributes to variability in the linear predictor also independently. When *M*_*x*_ is close to zero, although *V*_*x*_ is not, the linear term *x* has no direct effect on the linear predictor, but all variance attributed to *x* can be explained by other covariates. When negative, the linear term acts to supress variability in the linear predictor. All the previous cases imply that the unique variance partition *U*_*x*_ is smaller than *V*_*x*_. Lastly, when the variance partition *V*_*x*_ is equal to *M*_*x*_ even though *U*_*x*_ is smaller than *V*_*x*_, then the effect of the covariances averages out. To summarize, the difference between *V*_*x*_ and *U*_*x*_ serves as an indicator for the presence of covariances, and hence confounding, between the linear terms, while the difference between *V*_*x*_ and *M*_*x*_ shows whether the covariances increase or suppress the variability of the linear term or the linear predictor.

**Table 3.**
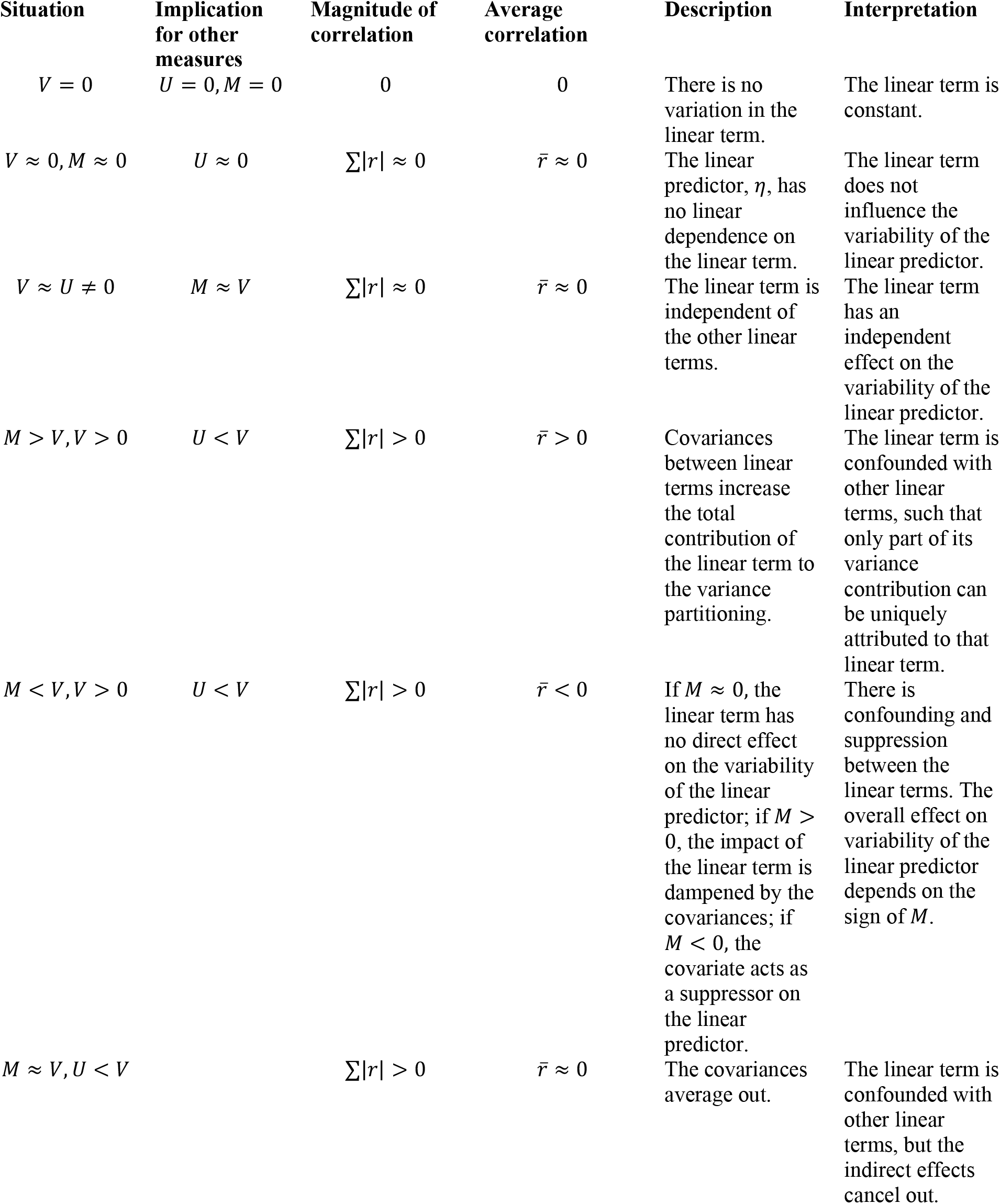
Joint interpretations of variance partitioning measures. *V* stands for the variance partition of a linear term or group of linear terms, *M* for its marginal variance partition and *U* for its unique variance partition. ∑|*r*| is the sum of absolute values of the correlation coefficients between the linear term and all other linear terms and 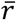 is the mean of the those correlation coefficients.

### 2.7 Variance partition measures revisited

For any variance partitioning measure there are many properties that would be desirable (Pratt 1987, Bring 1995, Nathans *et al*. 2012). We consider here only four of them (Table 2): the variance partitions should sum to one, their value should lie between zero and one, they should reflect the effect of the covariances and when the linear terms are uncorrelated they should all equal the variance partition *V*_*x*_. None of the measures presented here fulfill all of them unless the linear terms are uncorrelated (Table 2), which is why it can be useful to report variance partitions in terms of more than one measure (*cf*. Pratt 1987, Whittaker 1990, p. 130, Bring 1995). There are many other measures for variance partitioning that attempt to account for covariances to be found in the ecological and statistical literature. Some of the measures are simply reformulations of *V*_*x*_, *U*_*x*_, and *M*_*x*_. Others, which seemingly fulfill all of the properties mentioned above, have the unfortunate feature that they do not distinguish between direct and indirect variance contributions and hence do also not indicate the presence of confounding or suppression. Hence, they cannot be said to reflect the effect of covariances in an interpretable manner. Yet others, like commonality analysis, partition variance in to an exponentially increasing number of components as a function of linear terms or are not meaningfully defined for groups of linear terms of non-linear effects, which limit their application in practice. Thus we find that *U*_*x*_ and *M*_*x*_ are the most useful ones to study the effect of covariances as they are sufficient to answer elementary questions such as do correlations between the linear terms affect the variance partition (comparison of *U*_*x*_ and *V*_*x*_) and what is the direction and effect of the consequent covariances (comparison of *U*_*x*_, *M*_*x*_ and *V*_*x*_ together).

## 3 Variance partitioning in practice: Case study on the Glanville fritillary butterfly

We demonstrate the application of variance partitioning using a simple species distribution model to explore factors contributing to patch occupancy in a Glanville fritillary butterfly metapopulation in the Åland Islands, SW Finland (Fig. 1). We flesh out the methodological details of the variance partitioning by concisely explaining practical application of the theory to this case study. We demonstrate the analysis steps, present the results, and discuss them from the ecological interpretation point of view in this section. In Section 4 we provide thorough discussion on the findings from the methodological point of view.

**Figure 1.**
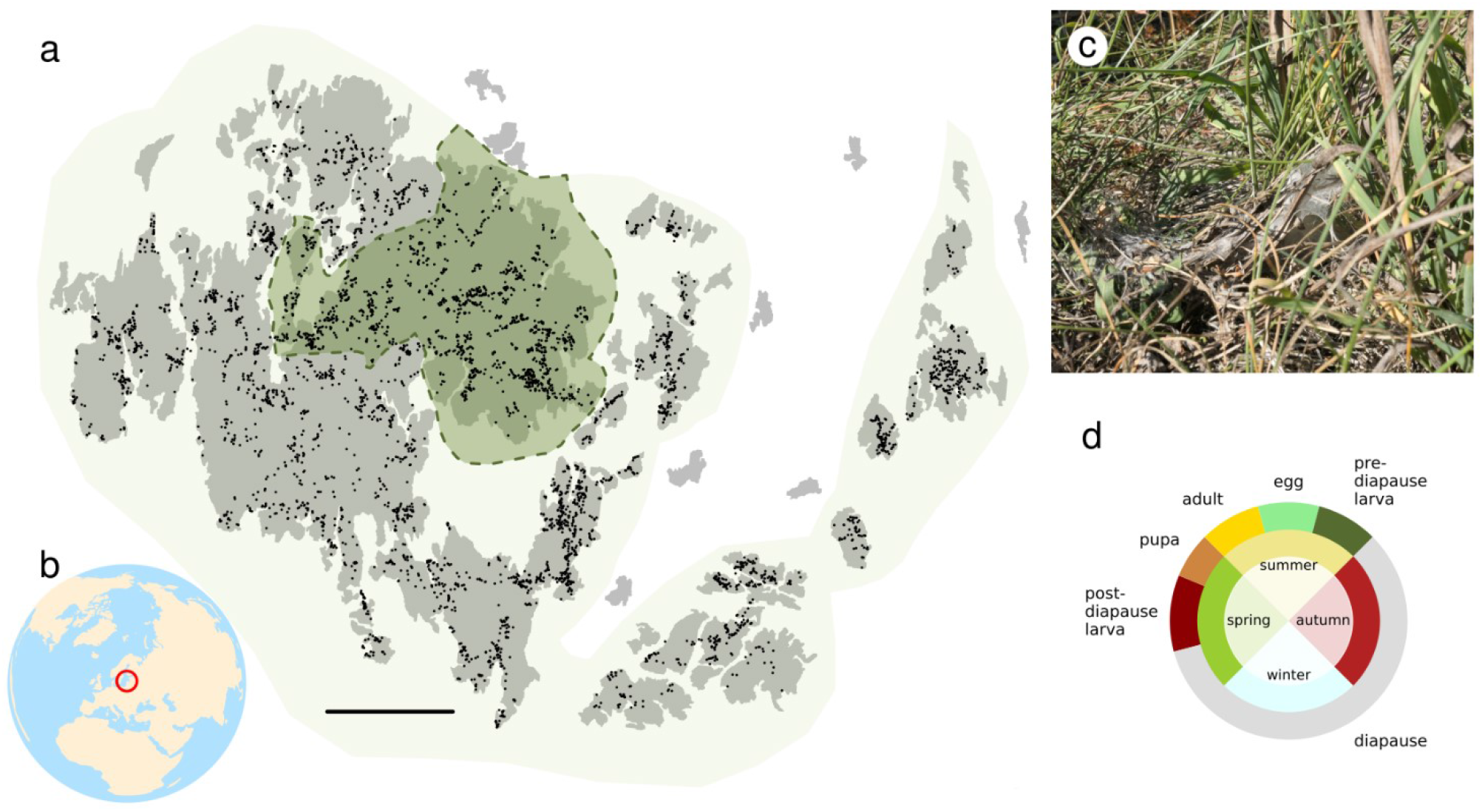
(**a**) Map of the Åland Islands study system. The black dots represent habitat patches suitable for *M. cinxia*. The darker enclosed region corresponds to the core area used for model inference, while the whole highlighted area corresponds to the “population” in our *population variance partitioning* measures. The solid black bar indicates 10 km in the map’s projected coordinate system ETRS-TM35FIN. Shoreline data from GSHHG version 2.3.7. (**b**) A projected view of the globe with the Åland Islands at its center (lat. 58.7 N lon. 21.7 E). The red circle encloses approximate location of Åland in the Baltic Sea. Land area data from Natural Earth. (**c**) Silken ‘winter nest’ spun by the larvae of *M. cinxia* on *Plantago lanceolata*. Photograph by Torsti Schulz. (**d**) Life cycle of *M. cinxia*.

### 3.1 Study system and species distribution model

The study system consists of approximately some four thousand potential habitat patches in which the butterflies’ larval host plants, *Plantago lanceolata* and *Veronica spicata*, occur. These habitat patches range from dry meadows, pastures, and rocky outcroppings to roadsides and yards. The butterfly, *Melitaea cinxia*, has a single generation at this northern latitude of its range; the adult butterflies emerge in early summer and the larvae, that live greagariously most of their lives, build silken nests for overwintering. The nests are easily distinguishable and are counted during an annual autumn survey. At the same time data about the habitat is gathered, such as measures of the abundance of the host plant and indicators of grazing (Ojanen *et al*. 2013).

The model and data are simplified from our previous study analysing the relationship between occupancy and abundance in this long-term ecological study system (Schulz *et al*. 2019, 2020). The data consists of over 60 000 observations from the potential habitat patches over a span of 19 years. In addition to the nest counts, the annual records include five measures of habitat quality (host plant *vegetation*, proportion of *dry vegetation, grazing intensity, grazing presence*, presence of *both host* plants), while the effects of metapopulation structure is represented by patch area and an annual estimate of population connectivity (Model supplement, Schulz *et al*. 2020). For our analyses, we only select a subset of the data available in order to mimic a scenario where only a part of the potential study domain is surveyed (see Fig. 1). This *core area* has the highest occupancy rates (average within 17.5 % *vs*. 14.5 % outside) and comprises *n*_*p*_ = 1259 of the total 4379 habitat patches in the data set. When we refer to unobserved patches or locations below we mean the patches outside this *core area*. In reality, the full data set contains observations also from those patches. Note that we use previous occupancy as a covariate in our model and it will be missing for cases when the corresponding habitat patch was not surveyed in the previous year. For demonstration purposes we ignore the problem and simply omit the observations for which there is no corresponding observation from the same patch in the previous year. We also exclude observations with no host plant present (n = 635) as it is really an indication that the patch is no longer habitat for the butterfly.

The occupancy state of the *i*th habitat patch observation is defined as *y*_*i*_ = *I*(*N*_*i*_ > 0), where *N*_*i*_ is the number of winter nests found. The vector of occupancy states 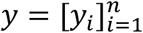, where *n* = 20086, collects all observations from the *n*_*p*_ = 1259 individual patches and *T* = 19 years. We model the occupancy state using logistic regression with the logit link-function such that E[*y*_*i*_] = logit^−1^(*η*_*i*_), where *η* is the linear predictor vector with elements

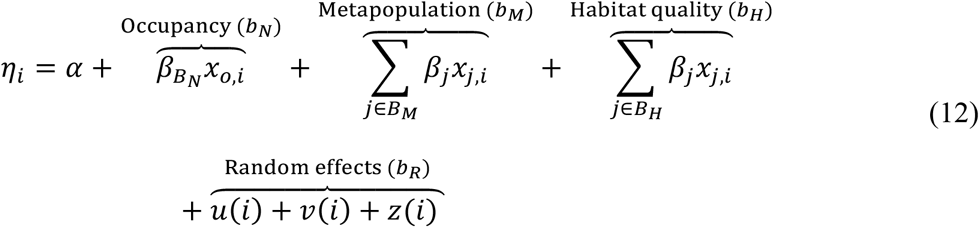

where *α* is the constant of regression, *x*_*o,i*_ *∈* {0,1} is the occupancy status of the patch in the previous year (0 representing unoccupied) and *occupancy, metapopulation*, and *habitat quality* index sets *B*_*N*_, *B*_*M*_ and *B*_*H*_ collect the “fixed effects” into covariate predictor groups (see section 2.3; note that the set *B*_*N*_ contains only one covariate index *x*_*o*_; Model supplement). The *random effects* are collected into an additional random effects group labeled *R*. We use this thematic grouping of the linear terms to study which general factors affect our study system rather than individual covariates or random effects. *u* and *v* are per patch and year *iid* random effects and *z* is a spatio-temporal random effect.

Throughout this running example, we assume that posterior inference is implemented so that we can produce random samples from the joint posterior distribution of the model parameters and random effects. This is the most common approach to implement Bayesian inference and allows using Monte Carlo techniques in calculating the variance partition measures. Hence, we assume the posterior distribution of model parameters are represented by vectors, such as [*α*_1_, …, *α*_*Q*_] where *Q* is the number of posterior samples. The posterior distributions of random effects are represented by matrices, such as [*u*_1_, …, *u*_*Q*_] where *u*_*q*_ is a column vector containing the *q*th posterior sample for all *n*_*P*_ observation locations. In our particular model, posterior inference was performed with R-INLA (Rue *et al*. 2009, Lindgren *et al*. 2011, Lindgren and Rue 2015; INLA version 21.02.23, R version 4.0.4) and is described in detail in the Model supplement where we also report the posterior distributions of the model’s parameters. The full code for the analyses is publicly available (Schulz *et al*. 2021). For code examples using a simpler model see the Programming supplement.

### 3.2 Sample variance partitioning for the regression model

We are interested in partitioning the variance of the linear predictor *η* into the groups of linear terms indicated by brackets above the model (Eq. (12)). That is, we partition the variance of linear predictor with respect to these four groups

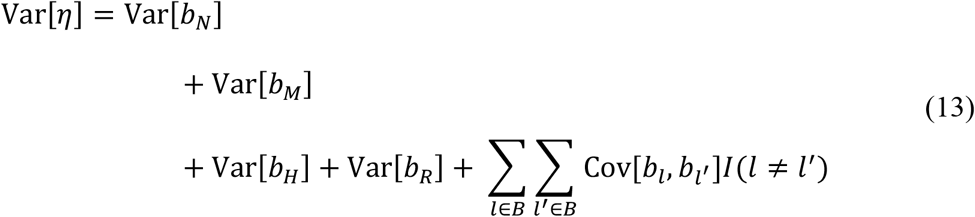

where *B* = {*N, M’, H, R*} collects the labels for the predictor groups. For example, 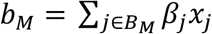, and the indicator function *I* ensures that the (diagonal) variances are not counted twice. The variance partition measures are then

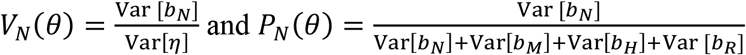

for occupancy and analogous for the other groups of linear terms (see also section 2.3 above).

To construct the posterior distribution of the sample variance partition we must first construct posterior samples of the predictor group effects. The samples for the fixed effects groups are specified as, 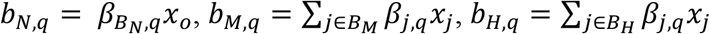 where *q* = 1, …, *Q* is the index of a Monte Carlo sample from the joint posterior distribution of all parameters and random effects and each *b* is a column vector with rows for each observation. A posterior sample for the random effects group is correspondingly constructed as *b*_*R,q*_ = *u*_*q*_ + *v*_*q*_ + *z*_*q*_. Thus the linear predictor is *η*_*q*_ = *b*_*N,q*_ + *b*_*M,q*_ + *b*_*H,q*_ + *b*_*R,q*_. Posterior samples for the variance partitions are then acquired as a function of the sample variance of the group of linear terms as

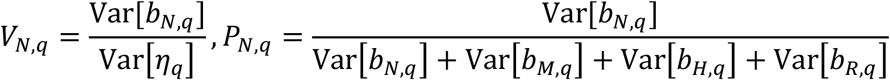

and 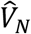 and 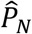 are then simply the mean over *V*_*N*,1_, … *V*_*N,Q*_ and *P*_*N*,1_, … *P*_*N,Q*_. The approach for the other predictor groups is identical.

The biased variance partitioning is calculated by first calculating posterior means of the predictor group effects: 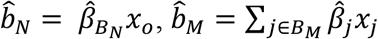, and 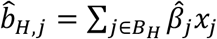 where 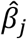 is the mean over *β*_*j*,1_, …, *β*_*j,Q*_. The posterior mean of the random effects group is constructed analogously. After this we plug in 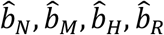 into the above equations to calculate 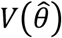 and 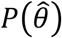.

The posterior distributions of all variance partitioning measures (e.g., *p*(*V*_*M*_|*D*)) exhibit considerable uncertainty (Fig. 2 (a)). The posterior means (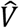 and 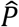) and posterior variances (Var[*V*|*D*] and Var[*P*|*D*]) of the variance partitioning measures summarize well the location and the spread of the variance partitions whereas the biased variance partitionings 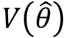 and 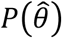. are considerably biased (Fig. 2 (a)). Moreover, compared to the posterior distribution of variance partitioning measures the biased variance partitionings (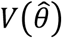 and 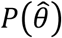) mistakenly suggest that the metapopulation predictor group would have higher contribution to total variance than the random effects predictor group. These differences in the summary measures illustrate the importance of reporting the variance partitioning based on the posterior distribution. The variance partition measures (*V*) are lower than the corresponding diagonal variance partitioning measures (*P*) indicating the presence of overall positive correlations among the predictor groups. The measures are qualitatively equivalent in that they preserve the ordering of the variances of the groups of linear terms, but the posterior distribution of their ratio, 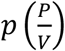, varies as a function of what proportion of variation in the linear predictor is due to the covariances.

**Figure 2.**
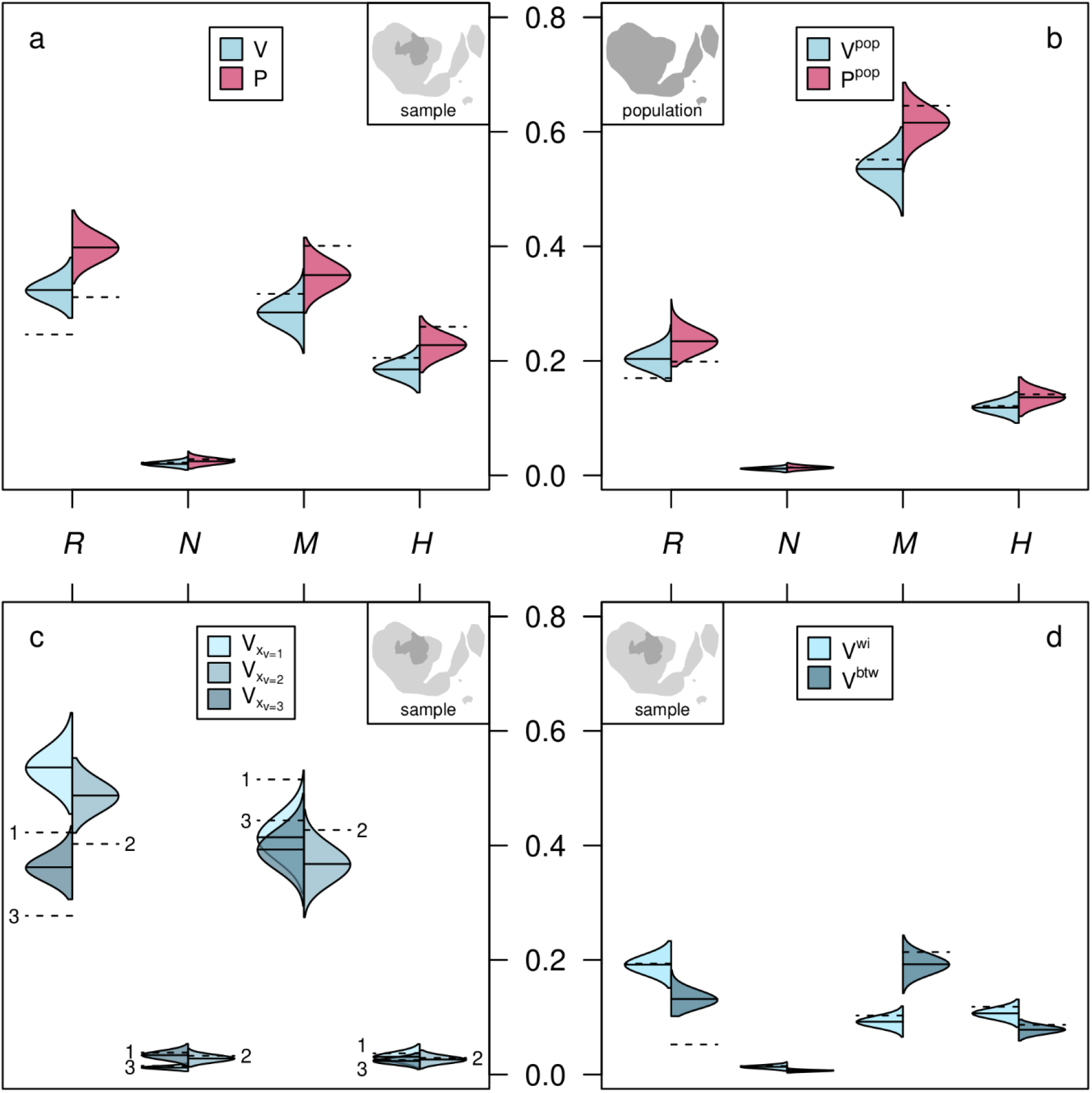
Variance partitioning of the linear predictor. The variance terms are grouped into four groups of linear terms: random effects (R) previous occupancy (N), metapopulation covariates (M), and habitat quality covariates (H). The violin plot distributions correspond to *p*(*V*|*D*) and *p*(*P*|*D*). Solid lines represent the posterior mean of 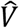 and 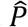 as the point estimate summary of the variance partition. Dashed lines represent the corresponding biased point estimate approximations with the posterior mean of the model parameters (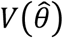 and 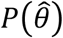). The inset maps show over which part of the study region the variance partition is calculated (see also Fig. 1). (**a**) Sample variance partitions and diagonal variance partitions for the core study area. (**b**) Variance partition including predictions to the survey areas not included in the model. (**c**) Sample variance partitioning conditional on the different levels of the host plant covariate (vegetation). (**d**) Sample variance partitions within and among patches.

The random effects, metapopulation covariates and the habitat quality covariates all contribute a large proportion to the total variation in occupancy in the core study area (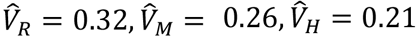, Fig. 2 (a)). The role of occupancy in the previous year in the variation of current occupancy is very low 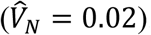. Partly this is due to expected correlations with the metapopulation covariates connectivity (*r* = 0.48) and area (*r* = 0.27) as well as host plant abundance (*r* = 0.22). Given the high rate of extinctions and colonizations previous occupancy alone is not a very good predictor of current occupancy in this system; and connectivity also is a predictor for colonizations while area as well as host plant abundance are good surrogates for population size, which does predict extinction risk (Schulz *et al*. 2020). In Section 3.6 we will take a more formal look at the role of these correlations.

### 3.3 Population variance partitioning

In our case study, model inference is performed only on a subset of the study region (Section 3.1, Fig. 2). However, we are interested in the relative importance of the groups of linear terms over the whole distribution range of Glanville fritillary in the Åland Islands. The locations, where data have been collected, correspond to the sample and all the locations within the distribution range of Glanville fritillary correspond to the population in our study (section 2.4). To calculate the *population variance partition*, we collect into 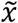 the covariate observations from the unsampled patches.

In order to construct samples from the posterior distribution of the population variance partitioning, we first construct posterior predictive samples for the predictor groups at unobserved patches. For fixed effects groups they are specified analogously to sample variance partitioning (section 3.2) as column vectors 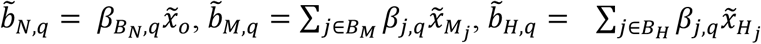 where *q* = 1, …, *Q* is the index of a Monte Carlo sample from the joint posterior distribution of all parameters and random effects. For the random effects’ predictor group 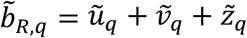 we need to construct the predicted random effects at the unobserved locations. In our case, the sample data cover the same years as the predictions, so the yearly random effects are simply 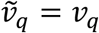. For the patch effects we must sample for each unobserved habitat patch new values from a univariate normal distribution with variance corresponding to the *q*th posterior sample of the variance parameter of the patch random effect. The case for the spatio-temporal random effect the approach is similar, but more complicated due to the covariance structure of the spatio-temporal random field which depends both on the hyperparameters defining the covariance matrix as well as the existing random effects *z*_*q*_ at the observed patches. For technical details see the Model supplement.

Once we have constructed the samples for predictor group predictors at the unobserved locations, for each sample we concatenate the unobserved predictor group column vector with the corresponding column vectors at the observed locations to form, 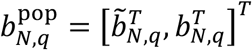 and equivalently for the other predictor groups. After this, we can proceed as before and calculate the *population variance partition* over the whole study region as

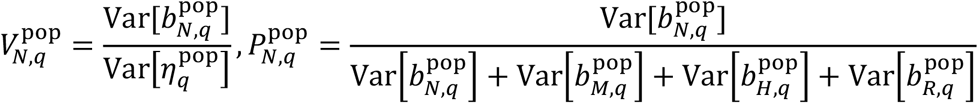

where 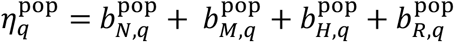 A variance partitioning calculated with only 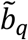, such as 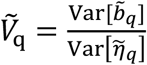 would correspond to *predictive variance partitioning* (section 2.4) over the unobserved locations. In principle, it would be straightforward to replace the predictive covariate values, 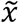 and 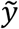, with some scenario-based covariates, such as covariates corresponding to climate or habitat change.

The population variance partitioning differs from the sample variance partitioning considerably in terms of the role of the metapopulation predictors (Fig. 2 (a) and (b)). Over the whole habitat range in the Åland Islands, it accounts for approximately half of the variation in the occupancy of the butterfly, with correspondingly reduced roles for the random effects and habitat quality 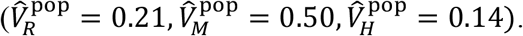.

This suggest that outside of the core study area connectivity and habitat patch size determine variation in occupancy as there is less variation in habitat quality (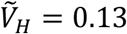 *vs*. 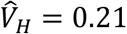). This is consistent with the fact that the core study area contains most of the more viable habitat of the butterfly in the Åland islands (Hanski *et al*. 2017). Also, the habitat patches outside of the core area tend to be more isolated accentuating the role of connectivity in predicting occupancy.

### 3.4 Variance partitioning conditional on host plant abundance

The amount of host plant vegetation within the habitat patch is known to be crucial for the specialist larvae of the butterfly (Opedal *et al*. 2020). We thus want to know what other factors may contribute to variation in occupancy after we have taken the amount of host plant vegetation into account. For this we partition variation conditional on the value of the host plant covariate (vegetation), *x*_*v*_, which conveniently for our demonstration is measured on an ordinal scale from 1 to 3 with 0 indicating absence of the host plant. We define the conditioning function *g*(*x*) = *x*_*V*_. The conditional variance partition is calculated just as the regular, but separately within the observations belonging to each value *c* = 1,2,3 of the vegetation covariate *x*_*v*_ (see section 2.5). The variance partition for occupancy conditional on the vegetation covariate is

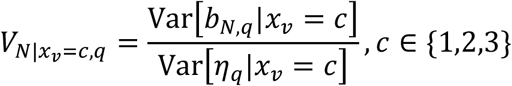

and is calculated equivalently for the other predictor groups.

Differences between the conditional variance would imply that the other covariates or the random effects covary with host plant abundance. Or more precisely, that the distribution of the covariates (and random effects) differs between the levels of host plant abundance. Here, we see mostly only small differences between the abundance classes except for the random effects, where less variation is associated with random effects in the highest abundance class (Fig. 2 (c)). The minor role of the habitat quality predictor group indicates that the predictor group’s variance contribution is mostly due to the conditioning variable host plant abundance. Again the posterior mean and variance estimates (e.g. 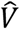 and Var[*V*|*D*]) summarize well the posterior distributions for variance partitioning. However, especially for the random effects, the biased point estimates greatly underestimate the conditional variance proportions.

### 3.5 Variance within and between patches

We go one step further and ask whether more variation in occupancy is due to differences between patches or due to annual variation within patches and which factors drive these differences (see also section 2.5). To do this, we define the conditioning function to be *g*(*x*) = *x*_*P*_ where *x*_*P*_ is an indicator vector of length *n* for *n*_*P*_ = 1259 patches, with *x*_*P,i*_ = *p*(*i*) above.

The variation within patches 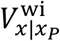 is a weighted average of their conditional variances, while the variance between patches 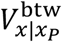 is the weighted variance of the patch means (averaged over time) of the predictor group effects. The weighing is necessary to allow for unbalanced group sizes, in this case differing numbers of observations per patch. A posterior sample from the within and between condition variance for occupancy are formed as

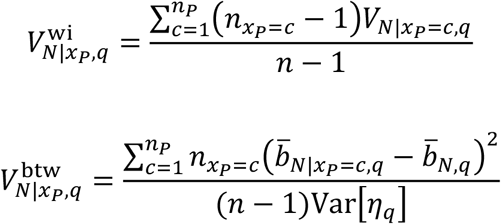

Here 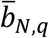 is the mean of occupancy over all observations, while 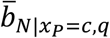 is the per-patch mean for occupancy at patch *c* both calculated with the *q*th Monte Carlo sample of parameters. That is 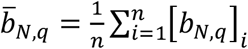 and, 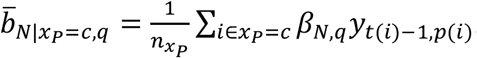. The calculation for the other predictor groups follow the same pattern.

When the variance contributions are separated for within and between habitat patches we see that the metapopulation covariates contribute more to differences in occupancy between patches than interannual variability within the patches (Fig. 2 (d)). This is to be expected as connectivity varies significantly between patches already simply due to varying isolation, though annual variation is also present (Hanski *et al*. 2017, Kahilainen *et al*. 2018). And as the data uses a single areal measurement for each patch, habitat area cannot contribute at all to within patch variation. For the random effects the situation is reversed and they contribute somewhat more to yearly variation in occupancy within the patches. The habitat quality covariates contribute almost equally to variation within and between patches. This dual role for habitat quality is consistent with the observation that in addition to habitat patches vastly differing in quality (Schulz *et al*. 2020) there is also significant annual variation in host plant availability within these patches (Opedal *et al*. 2020).

The biased point estimates are rather close to the actual variance partitions in all cases but one. The estimate for the random effects’ contribution to variation between patches is severely underestimated. A likely explanation lies in the fact that the posterior mean of the random effects is close to zero, which is both a function of the random effects’ priors having a mean of zero and an constraint for the random effects to sum to zero (Model supplement). As a consequence a large proportion of their variance contribution is due to the posterior variance, which is ignored by the biased point estimate. This is a general problem whenever the mean estimate is close to zero, but the uncertainty about the estimate is great, which almost by design is true of random effects, but can affect covariate terms as well.

### 3.6 Correlation between covariates and posterior correlation between parameters

To understand the effects of correlated covariates and random effects, we start by visualizing both the posterior distributions of the covariances between predictor groups and the correlations of the predictor group’s variance partitions (Fig. 3). The unique variance partition for the groups of linear effects is the proportion of variation due to the linear terms in that group, which cannot be expressed as a linear combination of linear terms belonging to other groups. The measure can calculated using the sample covariance matrix of the linear terms (Mathematical supplement 2.6.2):

**Figure 3.**
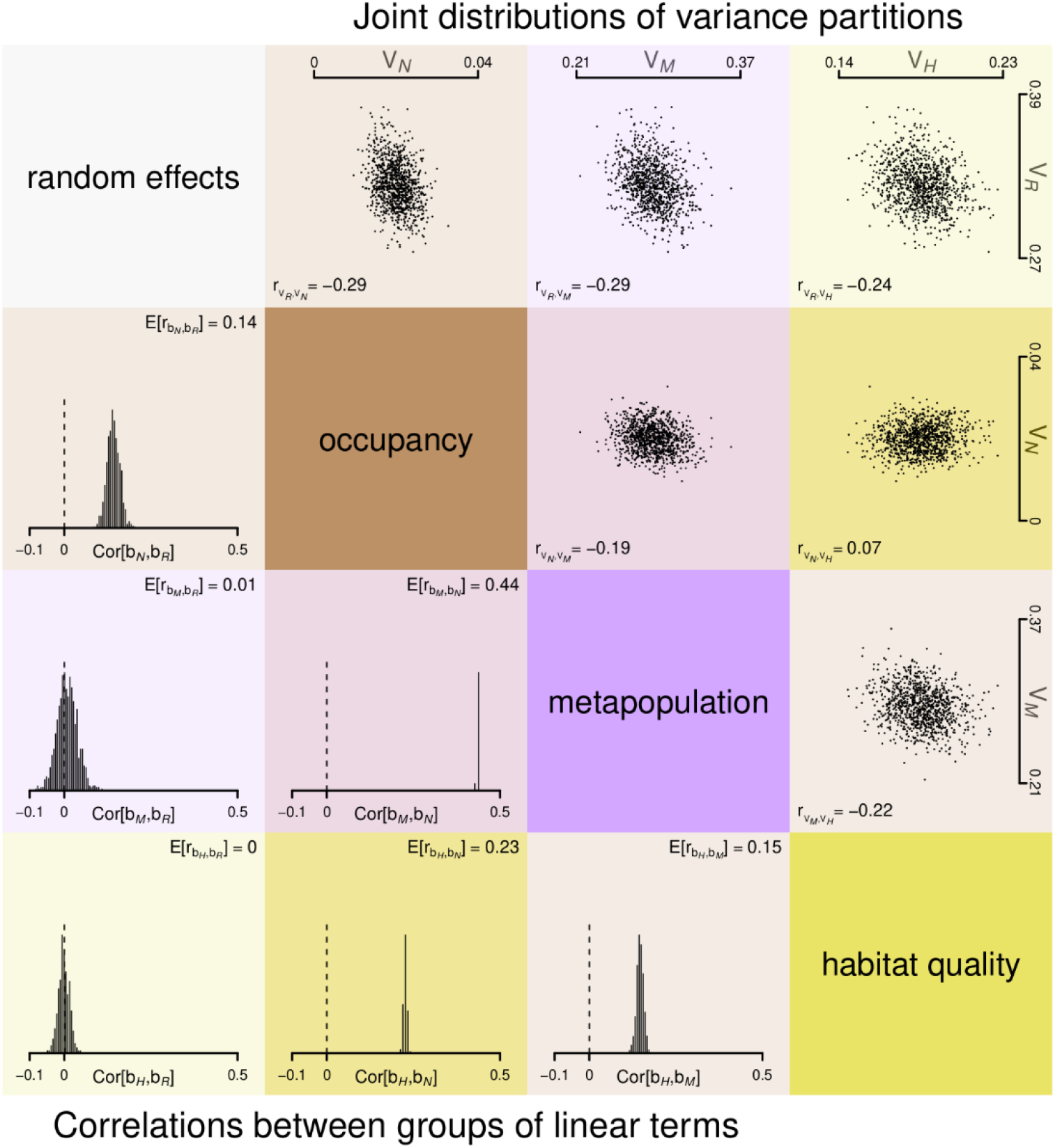
Posterior distributions of correlations between variance partitions and groups of linear terms for random effects (R) previous occupancy (N), metapopulation covariates (M), and habitat quality covariates (H). The histograms in the lower triangle represent the posterior marginal distributions of the correlations between the four groups of linear terms, *p*(Cor[*b*_*l*_, *b*_*l*′_]|*D*). Their posterior means are given in the upper right corner of each entry. The scatterplots in the upper triangle represent samples from the joint posterior distribution of pairs of variance partition measures 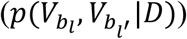. The posterior correlation between the variance partition measures is given in the lower left corner of each entry.

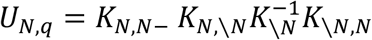

where *K* is the sample covariance matrix between the linear terms in the model, *N* indexes the rows or columns belonging to occupancy and \*N* all the linear terms in the other groups of linear terms. Note that in general *U*_*b*_ is the sum of the variances in the above equation, but that is omitted as the occupancy group consists of a single covariate.

The marginal variance partitions for groups of linear terms, *M*_*b*_, are calculated as the variance partition plus the covariance with the other groups:

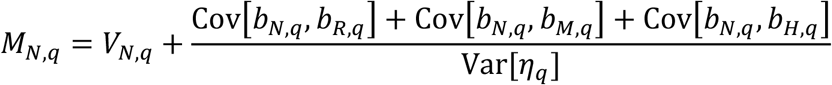

Some of the groups of linear terms with the high correlations between them have also high posterior correlations between their variance measures (see Fig. 3). But looking at correlations of pairs of linear term groups alone is not always sufficient to surmise the posterior correlations between their variance partitions. Rather, the correlations between the variance partitions posterior distributions are due to both the correlations among the covariates and the posterior correlations among the model parameters. Even if all covariates were mutually orthogonal, there could be posterior correlation between the variance partitions due to posterior correlations between model parameters. The effect of covariate correlations on the posterior correlation between two linear terms depends not only on the correlation between the corresponding covariates, but the size of their variance contributions and the correlations between all the other covariates. The resulting correlations are not always intuitive. As a stark example of this, the random effects and habitat quality covariates have a very low correlation, with the posterior distribution of covariances overlapping zero, yet their variance partitions have strong negative correlations (Cor[*V*_*R*_(*θ*), *V*_*H*_(*θ*)] = −0.33 by expectation). The same is true for the random effects and the metapopulation covariates (Cor[*V*_*R*_(*θ*), *V*_*M*_(*θ*)] = −0.23 by expectation). The posterior distribution of the random effects’ variance partition is negatively correlated with all the other variance partitions (Fig. 3) precisely because its variance contribution is mostly unique (Fig. 4). Any increase in its proportion of total variation will have to be offset by the other groups’ variance partitions. And the three groups of covariate linear terms are all correlated, but only the metapopulation and habitat quality variance partitions have a clear negative correlation (Fig. 3); the occupancy and the habitat quality linear term groups are clearly correlated (E[Cor[*b*_*N*_, *b*_*H*_]] = 0.23), but the correlation between their variance partitioning measures is close to zero.

**Figure 4.**
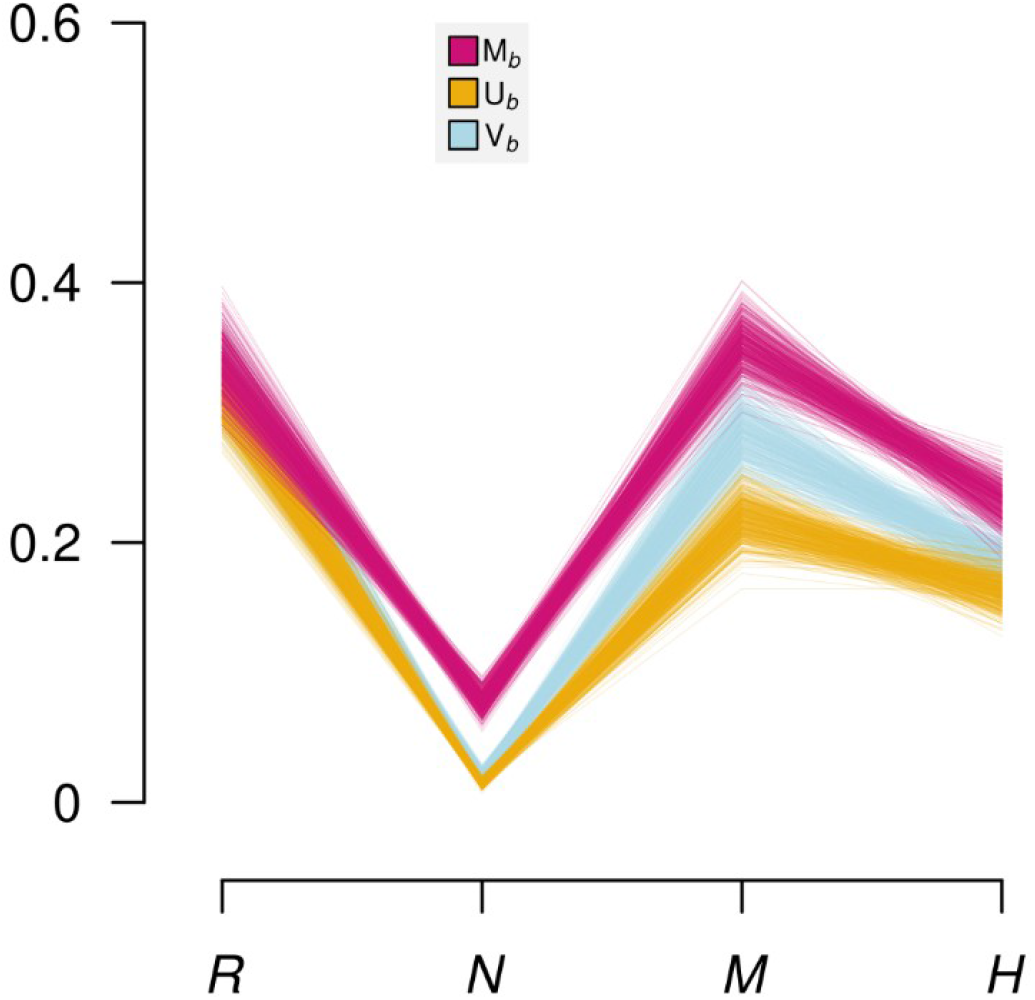
Parallel coordinates plot of the different measures of variance allocated to each group of linear terms. The groups are the random effects (R) previous occupancy (N), metapopulation covariates (M), and habitat quality covariates (H). The variance partitioning measures are the variance partition *V*_*b*_ in light blue, the unique variance partition *U*_*b*_ in orange and the marginal variance partition *M*_*b*_ in dark pink. Each line in the plot correspond to a sample from the (joint) posterior distribution of the measures.

The random’ effects unique variance partition *U*_*R*_ is approximately equal to its variance partition *V*_*R*_ and the marginal variance partition *M*_*R*_ is correspondingly only slightly higher than *V*_*R*_ (Fig. 4). This means there is only minor confounding between the random effects and the covariates, even though the posterior distribution of *V*_*R*_ is clearly correlated with the other variance partitions. All of the marginal variance partitions are higher than their corresponding variance partitions, which means that the covariances between the groups of linear terms are overall positive. The clearest example is occupancy, whose main contribution to variability is due to its covariances with the other linear terms. This is not unexpected as due to the high extinction and colonization rates in this system occupancy alone is not a generally reliable predictor of future occupancy states and as occupancy is highly correlated with habitat quality, patch area and connectivity. It is interesting to note that the metapopulation covariates and habitat quality covariate have approximately equal unique contributions, but the variance and marginal variance partition are both higher for the metapopulation covariates. A potential explanation is the higher correlation between occupancy and metapopulation covariates than occupancy and habitat quality covariates (Fig. 3).

## 4 Discussion

Variance partitioning in its many forms has been an important tool in ecology already for decades and we believe it continues to hold its central role in future as well. As often happens with highly influential methods, variance partitioning has evolved to many directions over the years. Some of these directions, however, have lost their connection to the original theory whereas others have improved and developed the core ideas significantly further. In this work, we have discussed the pros and cons of some of the modern versions of variance partitioning and proposed a few extensions to them that we see useful. Our work is not intended to be a review on the subject but we have taken a practical stance by pointing our criticism to popular approaches that we think are problematic while appraising approaches that we think should be more widely adopted in ecological research. We treat the subject from Bayesian perspective since it allows transparent and straightforward treatment of uncertainties.

The central idea in variance partitioning is to examine the relative importance of alternative ecological processes. As shown in our earlier presentation of the methods and exemplified in by our case study, the form of variance partitioning should change according to the question asked. For example, the sample variance partitioning for individual model components (the individual covariates in our case study) informs us about the relative roles of those covariates only (Section 2.2). However, distinct covariates often fall into broader sets of ecological processes, such as occupancy, metapopulation, habitat and random processes in our case study. If we are interested in the relative importance of these higher level processes in our study system, we should decrease the resolution of the analysis by using variance partitioning for the groups of linear terms formed by these upper level processes (Sections 2.3, 3.2). At the same time, the conditional variance partitioning and its extension to between and within condition variance partitioning increases the resolution of answers available by allowing the analyst to zoom into different compartments of the study system. That provides a method to assess what are the relative roles of the ecological processes in the subset of cases falling into a particular compartment, and whether the relative roles of the ecological processes are consistent between the compartments (low between condition variance) or not (high between condition variance) (Sections 2.5–2.6, 3.4–3.5).

Moreover, variance partitioning extends straightforwardly beyond the study system from where the data are collected. Here we demonstrated this by calculating the population variance partitioning for the larger system in whole from which we had (hypothetically) observed only a small fraction (section 3.3). We could analogously extend the idea to predictive variance partitioning analysis related to, for example, climate or land use change. These approaches to variance partitioning presented take a finite-population view of variation by looking at sample and population variance partitions for a specific study system. One essential question in ecology is, however, to distinguish between context dependency and generality. The variance partitioning can be turned to this question as well by implementing so called superpopulation view of variance (Gelman *et al*. 2014, pp. 201–202). For example, we could model spatially distinct study systems (such as, Glanville fritillary populations in Åland or UK) with a hierarchical Bayesian framework by assigning variance parameters to each study system and to a higher hierarchical level description of consistent environmental response across systems. The variance parameters could be interpreted as measures of variability across a larger domain than encompassed by any single study system. For specific questions such model could also make predictions for unobserved locations. In our context, this could for example involve the prediction of occupancy in populations elsewhere in the study species’ distribution. Superpopulation variance partitioning requires a more carefully thought out model structure and is more sensitive to choice of priors (Gelman 2005, 2006). Developing these methods further is beyond the scope of this work.

Our approach to variance partitioning is strictly model based, which has several consequences. All these analyses can be attained by a single model after a single parameter inference step. The model based approach allows us to coherently encode the ecological understanding into analysis. It also allows, when combined with Bayesian approach to parameter inference, transparent and theoretically justified treatment of uncertainty. These properties are clearly an advantage when our model explains the system well but it is also a reminder that we should not jump into variance partitioning before scrutinizing our model.

The treatment of uncertainty is one of the central themes in our work. In our experience, this is typically neglegted at many mutually connected levels of variance partitioning. Firstly, and most evidently, the variance partitioning summaries themselves are uncertain. This simple fact seems to be forgotten in most applications of variance partitioning even if authors had careful uncertainty treatment in other parts of their analysis. As shown in our results (Fig. 2), ignoring the uncertainty in variance partitioning by using simple parameter estimates may significantly change quantitative but also the qualitative conclusions from the study compared to treating the uncertainty correctly. Variance partitions should be treated as random variables and should be interpreted either by studying their distributions directly or by suppressing these distributions to well defined summary statistics such as posterior mean and variance or credible intervals. As shown in our case study, propagating the parameter uncertainty is rather simple in practice, as it can be done after model inference and only involves the same steps as calculating the variance partition in the first place. Hence, there is no reason to ignore the uncertainty in variance partitioning.

The above uncertainty considerations apply to the marginal distribution of variance partitions but another aspect of uncertainty is driven by the correlations between variance partitions. This is induced by correlations between model covariates and the correlations between model parameters in their posterior distribution. Even in the applied Bayesian variance partitioning studies that report uncertainties of marginal distributions, this aspect of uncertainty seems to have received barely any attention. For this reason we advocate explicitly reporting posterior correlations between variance partitions and linear terms as well as comparing marginal distribution of variance partitions to the unique variance partition and the marginal variance partition. These latter measures summarize the covariance relationships between the linear terms and, together with the basic variance partition, help interpreting possible confounding effects on the results.

When our research involves observational data, we will inevitably encounter situations where our covariates are correlated. The first step in approaching analyses of such data is acceptance. Correlations might be a nuisance from the analyst’s perspective, but they are also a real feature of the ecological system with implications for how the system works. In terms of partitioning variation, the resulting covariances between the linear terms are a feature not a deficiency in the method. They summarize the effects of the covariate distribution on variability and the limits of which effects can be identified independently. Hence we recommend not to remove covariates from an analysis simply because they are correlated (*cf*. Cox and Snell 1974, Morrisey and Ruxton 2018) as this only hides the *problem* and gives a false sense of certainty. We also do not recommend principal components analysis (PCA) as a general solution to correlated covariates. To remain meaningfully interpretable, the covariates included in PCA should be sufficiently similar. Moreover, in attempting to interpret the PCA one has not escaped the correlations after all, but moved the problem to the interpretation of the loading factors. And when working with non-linear or additive effects it does not seem sensible to use anything but the original covariates.

If the aim is to reduce the number of covariates to a smaller group of meaningful features, this can be achieved with grouping of the linear terms. Especially when interest is less in interpreting regression coefficients or prediction and more in looking at variance contributions, the problem of correlations is reduced by the grouping of the linear terms. When the correlation between covariates is strong enough that their independent effects cannot be estimated in a model, by grouping the linear terms one can still estimate their joint contribution to the model (*cf*. Mood 1971). Generally, we recommend only grouping thematically related covariates. Depending on the research question, the analyst might well be interested even in reporting variances for predictor groups consisting of rather dissimilar covariates. Either approach provides an alternative to excluding correlated covariates, a practice often arising from a misunderstanding about the effect and role of collinearity (Cox and Snell 1974, Morrisey and Ruxton 2018).

The simple fact that variances of sums equal sums of variances gets us surprisingly far. We like to think of the linear terms and their variances as the common currency of regression models. They are all on the same scale and can be meaningfully added together, which makes variance partitioning such a versatile approach — while we should not add apples and oranges, we can freely add growth from apples and oranges eaten to get total growth. Unlike working only directly with regression coefficients, partial regression coefficients, or especially non-linear effects, variance partitioning can deal with both individual predictors and groups of them. And the within–between condition comparison is not available from simple regression coefficients which also do not account for effect size *vs*. realized variability.

## Supporting information

Mathematical supplement

Programming supplement

Model supplement

## Acknowledgements

We wish to acknowledge Elina Numminen and Aapo Kahilainen for their critical comments on the manuscript. TS was funded by the Doctoral Programme in Wildlife Biology Fellowship (Univ. of Helsinki) and MS received funding from the European Research Council (Independent Starting grant no. 637412 ‘META-STRESS’) and JV from the Academy of Finland (grant 317255).

## Data availability

The code (Schulz *et al*. 2021) are available from Zenodo: [https://doi.org/10.5281/zenodo.5577956], The data (Schulz *et al*. 2019) are available from Dryad: [doi:10.5061/dryad.ksn02v707]

## Supplementary materials

Mathematical supplement

Programming supplement

Model supplement

