## Supplementary material for "Doors and corners of variance partitioning in statistical ecology": Mathematical supplement

### 1 Covariance structure of the Ricker model

We want to show that $\mathrm{Cov}[\beta_1 x_t, \beta_2 N_t] \neq 0$ when $\mathrm{Cov}[x_t, x_{t-1}] \neq 0$.

By definition $\mathrm{Cov}[\beta_1 x_t, \beta_2 N_t] = \mathrm{E}_{p(x_t,N_t)}[\beta_1 x_t \beta_2 N_t] - \mathrm{E}_{p(x_t)}[\beta_1 x_t]\mathrm{E}_{p(N_t)}[\beta_2 N_t]$ where $\mathrm{E}_{p(x_t,N_t)}[\beta_1 x_t \beta_2 N_t] = \mathrm{E}_{p(x_t,N_{t-1},x_{t-1},\epsilon_{t-1})}\left[\beta_1 x_t \beta_2 N_{t-1} e^{\alpha+\beta_1 x_{t-1}+\beta_2 N_{t-1}+\epsilon_{t-1}}\right]$. Since $\mathrm{Cov}[x_t, x_{t-1}] \neq 0$, it follows that $x_t$ and $N_{t-1}e^{\beta_1 x_{t-1}}$ are not independent and, thus, they are not uncorrelated in general. It follows that $\mathrm{E}_{p(x_t,N_t)}[\beta_1 x_t \beta_2 N_t] \neq \mathrm{E}_{p(x_t)}[\beta_1 x_t]\mathrm{E}_{p(N_t)}[\beta_2 N_t]$ and $\mathrm{Cov}[\beta_1 x_t, \beta_2 N_t] \neq 0$ in general.

### 2 Variance partitioning

#### 2.1 Variance of the linear predictor

##### 2.1.1 The linear predictor

The linear predictor, $\eta = (\eta_i) \in \mathbb{R}^n$, is a column vector formed as an unweighted linear combination of $d$ linear terms $\eta_i = \sum_{j=1}^{d} a_{i,j}$ or $\eta = A\mathbf{1}_d$ where $\mathbf{1}_d = (1_i) \in \{1\}^d$ is the summation vector of length $d$ and $A = (a_{i,j}) \in \mathbb{R}^{n\times d}$ is the matrix of linear terms with $a_{i,j} = f_j(x_i)$ or $a_{\cdot,j} = [a_{1,j}, \dots, a_{n,j}]' = f_j(x)$ for some matrix of covariates $x$. Note, for a linear regression model $a_{i,j} = x_{ij}\beta_j$. The elements of a vector $x_i, i \in \{1, \dots, n\}$, can be of any kind or dimension as long as the codomain of the function $f_j$ is either the real numbers $\mathbb{R}$ such that $a_{i,j} = f_j(x_i)$ or the vector space $\mathbb{R}^n$ such that $a_{\cdot,j} = f_j(x)$. The latter notation makes it more explicit that the values or rows of $a_{\cdot,j}$ can be correlated with each other as is the case with *e.g.*, spatial random effects.

##### 2.1.2 Variance of the linear predictor

The sample variance of the linear predictor is $\mathrm{Var}[\eta] = \frac{\sum_{i=1}^{n}(\eta_i - \bar{\eta})^2}{n-1}$ with the mean $\bar{\eta} = \frac{\sum_{i=1}^{n}\eta_i}{n}$. The variance can equivalently be written in terms of the matrix of linear terms (see also Whittaker 1990, pp. 121–124, Mulaik 2010, pp. 86–89):

$$\mathrm{Var}[\eta] = \mathrm{Var}[A\mathbf{1}_d]$$

$$\mathrm{Var}[\eta] = \frac{1}{n-1}\sum_{i=1}^{n}[(A\mathbf{1}_d)_i - \bar{\eta}\,]^2$$

$$\mathrm{Var}[\eta] = \frac{1}{n-1}\sum_{i=1}^{n}\left[(A\mathbf{1}_d)_i - \frac{1}{n}\mathbf{1}_n' A\mathbf{1}_d\right]^2$$

$$\mathrm{Var}[\eta] = \frac{1}{n-1}\left(A\mathbf{1}_d - \frac{1}{n}\mathbf{1}_n\mathbf{1}_n' A\mathbf{1}_d\right)'\left(A\mathbf{1}_d - \frac{1}{n}\mathbf{1}_n\mathbf{1}_n' A\mathbf{1}_d\right)$$

$$\mathrm{Var}[\eta] = \frac{1}{n-1}\left(\left(A - \frac{1}{n}\mathbf{1}_n\mathbf{1}_n' A\right)\mathbf{1}_d\right)'\left(\left(A - \frac{1}{n}\mathbf{1}_n\mathbf{1}_n' A\right)\mathbf{1}_d\right)$$

$$\mathrm{Var}[\eta] = \frac{1}{n-1}\mathbf{1}_d'\left(A - \frac{1}{n}\mathbf{1}_n\mathbf{1}_n' A\right)'\left(A - \frac{1}{n}\mathbf{1}_n\mathbf{1}_n' A\right)\mathbf{1}_d$$

$$\mathrm{Var}[\eta] = \frac{1}{n-1}\mathbf{1}_d'\left(A'A - \frac{1}{n^2}A'\mathbf{1}_n n\mathbf{1}_n' A\right)\mathbf{1}_d$$

$$\mathrm{Var}[\eta] = \frac{1}{n-1}\mathbf{1}_d' A' \left(I_n - \frac{1}{n}\mathbf{1}_n\mathbf{1}_n'\right) A\mathbf{1}_d$$
$$\mathrm{Var}[\eta] = \mathbf{1}_d{}' K_A \mathbf{1}_d$$

where $K_A$ is the sample covariance matrix of the columns of $A$, $K_A = (k_{j,j'}) \in \mathbb{R}^{d\times d}$, with $k_{j,j} \in \mathbb{R}_+$. Equivalently, $k_{j,j'} = \frac{\sum_{i=1}^{n}(a_{i,j}-\bar{a}_{\cdot,j})(a_{i,j'}-\bar{a}_{\cdot,j'})}{n-1} = \mathrm{Cov}[a_{\cdot,j}, a_{\cdot,j'}]$. Here, the means are as before $\bar{a}_{\cdot,j} = \frac{\sum_{i=1}^{n} a_{i,j}}{n}$. Hence, the variance of the linear predictor can also be written in terms of the variances and covariances of the individual linear terms as $\mathrm{Var}[\eta] = \sum_{j=1}^{d}\sum_{j'=1}^{d}\mathrm{Var}[a_{\cdot,j}, a_{\cdot,j'}]$.

#### 2.1.3 Correlations between the linear terms

To characterize the linear dependence between the linear terms we can also use their correlation, $\mathrm{Cor}[a_{\cdot,j}, a_{\cdot,j'}] = \frac{\mathrm{Cov}[a_{\cdot,j}, a_{\cdot,j'}]}{\sqrt{\mathrm{Var}[a_{\cdot,j}]}\sqrt{\mathrm{Var}[a_{\cdot,j'}]}}$. The sample correlation matrix of the linear terms is

$$R_A = \mathrm{diag}(K_A)^{-1/2} K_A \mathrm{diag}(K_A)^{-1/2}$$

so that $\mathrm{Cor}[a_{\cdot,j}, a_{\cdot,j'}] = r_{j,j'}$.

### 2.2 VARIANCE PROPORTIONS FOR PARTITIONING VARIATION

#### 2.2.1 Normalized variance partition

We define variance proportion or *variance partition* of the $j$th linear term as $V_j = \frac{\mathrm{Var}[a_{\cdot,j}]}{\mathrm{Var}[\eta]}$, that is $V_j$ is the variance of the $j$th linear term divided by the variance of the linear predictor. In matrix notation the variance partition is $V_j = \frac{K_{A\,j,j}}{\mathbf{1}' K_A \mathbf{1}}$. For the $V_j$s it holds that $\sum(V) = \sum_{j=1}^{d} V_j > 0$ and the difference $1 - \sum(V)$ equals the sum of the normalized covariances, $\sum_{j}^{d-1}\sum_{j'=j+1}^{d} 2\frac{K_{A\,j,j'}}{\mathrm{Var}[\eta]}$. When $\sum(V) < 1$ the sum of the covariances is positive, and it is negative when $\sum(V) > 1$.

#### 2.2.2 Diagonal variance partition

We denote by $P_j = \frac{\mathrm{Var}[a_{\cdot,j}]}{\sum_{j'=1}^{d}\mathrm{Var}[a_{\cdot,j'}]}$ the *diagonal variance partition*, where the diagonality comes from the normalizing denominator in matrix notation as $P_j = \frac{K_{A\,j,j}}{\mathrm{tr}(K_A)}$. The diagonal variance partition has the attractive quality $\sum_{j=1}^{d} P_j = 1$, but consequently carries no information about covariances between the linear terms. The variance in the denominator also equals the “total variation” in PCA (Legendre and Legendre 2012, pp. 165–171, 429–434, Jolliffe and Cadima 2016).

### 2.3 GROUPING LINEAR TERMS FOR VARIANCE PARTITIONING

If we partition the linear terms to a set of $m$ disjoint groups, so that each linear term, $a_{\cdot,j}$, belongs to exactly one group, then we can calculate the variance partition for each of these groups. Let $B \in \{0,1\}^{d\times m}$, be a binary matrix with $B_{j,l} = 1$ if the $j$th linear term belongs to group $l$ and 0 otherwise. As each linear term must only belong to one group it holds that $\mathbf{1}_d = B\mathbf{1}_m$. Now $AB = (b_{i,l}) \in R^{n\times m}$ is a matrix of grouped linear terms where $b_l = (AB)_{\cdot,l}$. We can also write $b_l = \sum_{j\in B_l} a_{\cdot,j}$ where $j \in B_l$ is a slight abuse of notation and denotes all the indices of all linear terms belonging to the $l$th group. The linear predictor can be expressed as $\eta = AB\mathbf{1}_m$ and its variance as (see also Whittaker 1990, 122–123, Mulaik 2010, pp. 91–92)

$$\mathrm{Var}[\eta] = \mathrm{Var}[AB\mathbf{1}_m]$$
$$\mathrm{Var}[\eta] = \mathbf{1}_m'\mathrm{Var}[AB]\mathbf{1}_m$$
$$\mathrm{Var}[\eta] = \mathbf{1}_m'B'\mathrm{Var}[A]B\mathbf{1}_m$$
$$\mathrm{Var}[\eta] = \mathbf{1}_m'B'K_AB\mathbf{1}_m$$

Now the variance partition for the $l$th group is $V_l = \frac{(B'K_AB)_{l,l}}{\mathrm{Var}[\eta]} = \frac{\mathrm{Var}[b_{\cdot,l}]}{\mathrm{Var}[\eta]} = \frac{\sum_{j\in l}\sum_{j'\in l}\mathrm{Var}[a_{\cdot,j},a_{\cdot,j'}]}{\mathrm{Var}[\eta]}$. The corresponding diagonal variance partition is $P_l = \frac{(B'K_AB)_{l,l}}{\mathrm{tr}(B'K_AB)} = \frac{\mathrm{Var}[b_{\cdot,l}]}{\sum_{l'=1}^{m}\mathrm{Var}[b_{\cdot,l'}]}$

### 2.4 CONDITIONAL VARIANCE PARTITIONING

#### 2.4.1 Notation for conditioning

We can partition the variance of the linear predictor into variation within and between groups of observations by a surjection $g(x)$ which assigns each of the $n$ observations to one of $n_G = |G|$ groups. The set of groups in the division is also called a set partition. Very commonly the function $g(x)$ could simply assign each observation to different values of a categorical variable. With $c \in G$, let $g(x_i)$ denote the group membership of the $i$th observation while we (ab)use $g = c$ to denote the set of indices of observations for which $g(x_i) = c$, that is, $g = c$ is shorthand for the set $\{i: i \in \{1, \dots, n\} \wedge g(x_i) = c\}$, where $c \in G$. Then $n_{g=c} = |g = c|$ is the number of observations in the group $c$.

#### 2.4.2 Conditional variance partitioning

The mean of the group $c$ is $\bar{\eta}_{g=c} = \frac{\sum_{i\in g=c}\eta_i}{n_{g=c}}$ and the conditional sample variance of group $c$ is $\mathrm{Var}[\eta_{g=c}] = \frac{\sum_{i\in g=c}(\eta_i - \bar{\eta}_{g=c})^2}{n_{g=c}-1}$. The equivalent partition for the linear terms is $V_{j|g=c} = \frac{\mathrm{Var}[a_{g=c,j}]}{\mathrm{Var}[\eta_{g=c}]}$ where $\mathrm{Var}[a_{g=c,j}] = \frac{\sum_{i\in g=c}(a_{i,j} - \bar{a}_{g=c,j})^2}{n_{g=c}-1}$ and $\bar{a}_{g=c,j} = \frac{\sum_{i\in g=c}a_{i,j}}{n_{g=c}}$. See also Whittaker (1990, pp. 124–125).

### 2.5 VARIATION WITHIN AND BETWEEN CONDITIONS

Let $\eta_{\bar{c}} = (\eta_{\bar{c},i}) \in \mathbb{R}^n$ be a vector of per observation group means such that $\eta_{\bar{c},i} = \bar{\eta}_{g=g(x_i)}$. That is, the $i$th element of $\eta_{\bar{c}}$ equals the mean of all the elements of linear predictor which belong to the same group as the $i$th element. Then we can write the linear predictor as the sum of a group mean and within group offsets.

$$\eta = \overbrace{(\eta_{\bar{c}})}^{\substack{\text{Condition}\\\text{mean}}} + \overbrace{(\eta - \eta_{\bar{c}})}^{\substack{\text{Within}\\\text{condition}\\\text{offset}}}$$

Now the variance of the linear predictor can also be split into variation between groups, $\mathrm{Var}[\eta_{\bar{c}}]$, and within groups, $\mathrm{Var}[\eta - \eta_{\bar{c}}]$:

$$\mathrm{Var}[\eta] = \mathrm{Var}[\eta_{\bar{c}}] + \mathrm{Var}[\eta - \eta_{\bar{c}}] + 2\mathrm{Cov}[\eta_{\bar{c}}, \eta - \eta_{\bar{c}}] = \underbrace{\mathrm{Var}[\eta_{\bar{c}}]}_{\substack{\text{Between-}\\\text{conditions}\\\text{variance}}} + \underbrace{\mathrm{Var}[\eta - \eta_{\bar{c}}]}_{\substack{\text{Within-}\\\text{conditions}\\\text{variance}}}$$

where the last equality arises because the covariance of the within and between condition terms, $\mathrm{Cov}[\eta_{\bar{c}}, \eta - \eta_{\bar{c}}] = 0$. We derive this result and the explicit equations for the between and within condition variances in the sections 2.5.1–2.5.3.

#### 2.5.1 Covariance between the centered values and groupwise means

Naively the partition is $\mathrm{Var}[\eta] = \mathrm{Var}[\eta_{\bar{c}}] + \mathrm{Var}[\eta - \eta_{\bar{c}}] + 2\mathrm{Cov}[\eta_{\bar{c}}, \eta - \eta_{\bar{c}}]$, hence we show that the covariance term of the within condition offsets and between condition means vanishes (*cf*. Whittaker 1990, pp. 124–125, Casella and Berger 2002, p. 537).

$$\mathrm{Cov}[\eta_{\bar{c}}, \eta - \eta_{\bar{c}}] = \frac{1}{n-1}\sum_{i=1}^{n}\left[\eta_{\bar{c},i} - \frac{1}{n}\sum \eta_{\bar{c},i}\right]\left[(\eta_i - \eta_{\bar{c},i}) - \bar{\eta} - \frac{1}{n}\sum \eta_{\bar{c},i}\right]$$

The mean of the within condition means is the grand mean, $\frac{1}{n}\sum \eta_{\bar{c},i} = \bar{\eta}$, and the mean of the within condition offsets is zero, that is $\bar{\eta} - \frac{1}{n}\sum \eta_{\bar{c},i} = 0$. Hence, the covariance is

$$\mathrm{Cov}[\eta_{\bar{c}}, \eta - \eta_{\bar{c}}] = \frac{1}{n-1}\sum_{i=1}^{n}[\eta_{\bar{c},i} - \bar{\eta}][(\eta_i - \eta_{\bar{c},i})]$$

We can replace the summing over the observations by the nested sum of the observations in each group and expand the product:

$$\mathrm{Cov}[\eta_{\bar{c}}, \eta - \eta_{\bar{c}}] = \frac{1}{n-1}\sum_{c\in G}\sum_{i\in g=c}[\eta_{\bar{c},i} - \bar{\eta}][(\eta_i - \eta_{\bar{c},i})]$$
$$\mathrm{Cov}[\eta_{\bar{c}}, \eta - \eta_{\bar{c}}] = \frac{1}{n-1}\sum_{c\in G}\sum_{i\in g=c}[\bar{\eta}_{g=c} - \bar{\eta}][(\eta_i - \bar{\eta}_{g=c})]$$
$$\mathrm{Cov}[\eta_{\bar{c}}, \eta - \eta_{\bar{c}}] = \frac{1}{n-1}\sum_{c\in G}\sum_{i\in g=c}[-\bar{\eta}_{g=c}^2 + \bar{\eta}_{g=c}\bar{\eta} + \eta_i\bar{\eta}_{g=c} - \bar{\eta}\eta_i]$$
$$\mathrm{Cov}[\eta_{\bar{c}}, \eta - \eta_{\bar{c}}] = \frac{1}{n-1}\left(-\sum_{c\in G}\sum_{i\in g=c}\bar{\eta}_{g=c}^2 + \sum_{c\in G}\sum_{i\in g=c}\bar{\eta}_{g=c}\bar{\eta} + \sum_{c\in G}\sum_{i\in g=c}\eta_i\bar{\eta}_{g=c} - \sum_{c\in G}\sum_{i\in g=c}\bar{\eta}\eta_i\right)$$

We move the constant scalars from the innermost sum to the outer sum over the groups:

$$\mathrm{Cov}[\eta_{\bar{c}}, \eta - \eta_{\bar{c}}] = \frac{1}{n-1}\left(-\sum_{c\in G}\bar{\eta}_{g=c}^2\sum_{i\in g=c}1 + \sum_{c\in G}\bar{\eta}_{g=c}\bar{\eta}\sum_{i\in g=c}1 + \sum_{c\in G}\bar{\eta}_{g=c}\sum_{i\in g=c}\eta_i - \sum_{c\in G}\bar{\eta}\sum_{i\in g=c}\eta_i\right)$$

We replace the inner sum operations by the equivalent expressions: $\sum_{i\in g=c} 1 = n_{g=c}$ and $\sum_{i\in g=c}\eta_i = n_{g=c}\bar{\eta}_{g=c}$:

$$\mathrm{Cov}[\eta_{\bar{c}}, \eta - \eta_{\bar{c}}] = \frac{1}{n-1}\left(-\sum_{c\in G}\bar{\eta}_{g=c}^2 n_{g=c} + \sum_{c\in G}\bar{\eta}_{g=c}\bar{\eta}n_{g=c} + \sum_{c\in G}\bar{\eta}_{g=c}n_{g=c}\bar{\eta}_{g=c} - \sum_{c\in G}\bar{\eta}n_{g=c}\bar{\eta}_{g=c}\right)$$

Rearranging the terms makes evident that the sums cancel out and the covariance is zero:

$$\mathrm{Cov}[\eta_{\bar{c}}, \eta - \eta_{\bar{c}}] = 0$$

#### 2.5.2 Between-condition variance

$$\mathrm{Var}[\eta_{\bar{c}}] = \frac{1}{n-1}\sum_{i}^{n}\left[\eta_{\bar{c},i} - \frac{\sum_i^n \eta_{\bar{c},i}}{n}\right]^2$$

$$\mathrm{Var}[\eta_{\bar{c}}] = \frac{1}{n-1}\sum_{i}^{n}\left[\eta_{\bar{c},i} - \bar{\eta}\right]^2$$

$$\mathrm{Var}[\eta_{\bar{c}}] = \frac{1}{n-1}\sum_{c\in G}\sum_{i\in g=c}\left[\eta_{\bar{c},i} - \bar{\eta}\right]^2$$

$$\mathrm{Var}[\eta_{\bar{c}}] = \frac{1}{n-1}\sum_{c\in G}\sum_{i\in g=c}\left[\bar{\eta}_{g=c} - \bar{\eta}\right]^2$$

$$\mathrm{Var}[\eta_{\bar{c}}] = \frac{1}{n-1}\sum_{c\in G} n_{g=c}\left[\bar{\eta}_{g=c} - \bar{\eta}\right]^2$$

$$\mathrm{Var}[\eta_{\bar{c}}] = \frac{\sum_{c\in G} n_{g=c}\left[\bar{\eta}_{g=c} - \bar{\eta}\right]^2}{n-1}$$

#### 2.5.3 Within-condition variance

$$\mathrm{Var}[\eta - \eta_{\bar{c}}] = \frac{1}{n-1}\sum_{i=1}^{n}\left[\left(\eta_i - \eta_{\bar{c},i}\right) - \left(\frac{\sum_{i=1}^{n}\left(\eta_i - \eta_{\bar{c},i}\right)}{n}\right)\right]^2$$

$$\mathrm{Var}[\eta - \eta_{\bar{c}}] = \frac{1}{n-1}\sum_{i=1}^{n}\left[\eta_i - \eta_{\bar{c},i}\right]^2$$

$$\mathrm{Var}[\eta - \eta_{\bar{c}}] = \frac{1}{n-1}\sum_{c\in G}\sum_{i\in g=c}\left[\eta_i - \eta_{\bar{c},i}\right]^2$$

$$\mathrm{Var}[\eta - \eta_{\bar{c}}] = \frac{1}{n-1}\sum_{c\in G}\sum_{i\in g=c}\left[\eta_i - \bar{\eta}_{g=c}\right]^2$$

$$\mathrm{Var}[\eta - \eta_{\bar{c}}] = \frac{\sum_{c\in G}\sum_{i\in g=c}\left[\eta_i - \bar{\eta}_{g=c}\right]^2}{n-1}$$

$$\mathrm{Var}[\eta - \eta_{\bar{c}}] = \frac{\sum_{c\in G}\left(n_{c=g} - 1\right)\frac{\sum_{i\in g=c}\left[\eta_i - \bar{\eta}_{g=c}\right]^2}{\left(n_{c=g} - 1\right)}}{n-1}$$

$$\mathrm{Var}[\eta - \eta_{\bar{c}}] = \frac{\sum_{c\in G}\left(n_{c=g} - 1\right)\mathrm{Var}\left[\eta_{g=c}\right]}{n-1}$$

Hence, the within condition variance is calculated from the conditional sums of squares or equivalently the weighted average of the conditional variances, where the weights are $\left(n_{c=g} - 1\right)$, but the average is taken over $n - 1$, unlike the usual definition of weighted averages or pooled variance as used in ANOVA (*cf.* Casella and Berger 2002, p. 528).

#### 2.5.4 Partitioning the linear terms

This partitioning to within and between conditions applies also to the linear terms and their covariances. With the matrix of per-observation linear term condition means $A_{\bar{c}} = \left(a_{\bar{c},i,j}\right) \in \mathbb{R}^n \times d$ where $a_{\bar{c},i,j} = \bar{a}_{g=g(i),j}$ we can partition the sample covariance matrix of the linear terms

$$\mathrm{Var}[A] = \mathrm{Var}[A_{\bar{c}}] + \mathrm{Var}[A - A_{\bar{c}}]$$

$$K_A = K_{A_{\bar{c}}} + \left(K_A - K_{A_{\bar{c}}}\right)$$

From their diagonal entries, we have the between conditions variance partition $V_j^{\text{btw}} = \frac{\left(K_{A_{\bar{c}}}\right)_{j,j}}{\text{Var}[\eta]} = \frac{\text{Var}[a_{\bar{c},\cdot,j}]}{\text{Var}[\eta]}$ and the within conditions variance partition $V_j^{\text{wi}} = \frac{\left(K_A - K_{A_{\bar{c}}}\right)_{j,j}}{\text{Var}[\eta]} = \frac{\text{Var}[a_{\cdot,j} - a_{\bar{c},\cdot,j}]}{\text{Var}[\eta]}$ of the linear terms.

The proof that the cross-covariances vanish is analogous to that of the covariances for the variance of the linear predictor (see also section 0).

#### 2.5.5 Partitioning variation within and between conditions using matrix notation

To avoid the sums-over-sets notation used above, we define the set partition via a binary matrix $H = \left(h_{i,c}\right) \in \{0,1\}^{n \times n_G}$ with $h_{i,c} = I(g(x_i) = c)$. This implies that each observation $i$ is assigned to only one group or condition $c \in G$, or equivalently that $\mathbf{1}_n = H\mathbf{1}_{n_G}$. Then $H'\mathbf{1}_n$ is a vector of length $n_G$ and its entries, $n_{g=c}$ are the numbers of observations in each group $c$. Let $\bar{A}_C = \text{diag}(H'\mathbf{1}_n)^{-1}H'A$ be a $n_G \times d$ matrix of group-wise linear term means with its elements equal to $\bar{a}_{g=c,j}$. Then $A_{\bar{c}} = H\bar{A}_C$ is the matrix of per observation group-means of the linear terms. Now we can proceed with partitioning the variance of $A$

$$\text{Var}[A] = \text{Var}[A_{\bar{c}}] + \text{Var}[A - A_{\bar{c}}] + 2\text{Cov}[A_{\bar{c}}, A - A_{\bar{c}}] = \text{Var}[A_{\bar{c}}] + \text{Var}[A - A_{\bar{c}}]$$

Note that the cross-covariance is

$$\text{Cov}[A_{\bar{c}}, A - A_{\bar{c}}] = \frac{1}{n-1}\left(A'_{\bar{c}}(A - A_{\bar{c}}) - \frac{1}{n}A'_{\bar{c}}\mathbf{1}_n\mathbf{1}'_n(A - A_{\bar{c}})\right) = 0$$

where the last equality can be shown analogously to the proof in Section 2.5.1. The between conditions covariance is

$$\text{Var}[A_{\bar{c}}] = \frac{1}{n-1}\left(A'_{\bar{c}}A_{\bar{c}} - \frac{1}{n}A'_{\bar{c}}\mathbf{1}_n\mathbf{1}'_nA_{\bar{c}}\right)$$

The within conditions covariance is

$$\text{Var}[\,A - A_{\bar{c}}] = \frac{1}{n-1}\left((A - A_{\bar{c}})'(A - A_{\bar{c}}) - \frac{1}{n}(A - A_{\bar{c}})'\mathbf{1}_n\mathbf{1}'_n(A - A_{\bar{c}})\right)$$

#### 2.5.6 Partitioning of grouped terms

Given that $K_A = K_{A_{\bar{c}}} + \left(K_A - K_{A_{\bar{c}}}\right)$ and $K_{AB} = B'K_AB$ (see section 2.3), we can write the grouped covariance matrices as

$$B'K_AB = B'\left(K_{A_{\bar{c}}} + \left(K_A - K_{A_{\bar{c}}}\right)\right)B$$
$$B'K_AB = B'K_{A_{\bar{c}}}B + B'\left(K_A - K_{A_{\bar{c}}}\right)B$$

and express the between grouped conditions variance partition as $V_l^{\text{btw}} = \frac{\left(B'K_{A_{\bar{c}}}B\right)_{l,l}}{\text{Var}[\eta]}$ and the grouped within conditions variance partition as $V_l^{\text{wi}} = \frac{\left(B'\left(K_A - K_{A_{\bar{c}}}\right)B\right)_{l,l}}{\text{Var}[\eta]}$ for the linear terms.

### 2.6 ALTERNATIVE MEASURES OF VARIATION

#### 2.6.1 Unique variance partition

We define the the *unique variance* partition as the semi-partial correlation coefficient between a linear predictor and a linear term. It corresponds to the variance of the least-squares prediction of the linear predictor from the residuals of a linear term regressed on all the other linear terms

$$U_j = \frac{\mathrm{Var}\left[\hat{\eta}\left(a_{\cdot,j} - \hat{a}_j(A_{\setminus j})\right)\right]}{\mathrm{Var}[\eta]}$$

where $A_{\setminus j}$ is the matrix of linear terms excluding the $j$th linear term, the hat-notation denotes the linear least squares predictor such that $\hat{a}_j(A_{\setminus j})$ is the least squares prediction for $a_{\cdot,j}$ conditional on the linear terms in $A_{\setminus j}$. Hence, $\hat{\eta}\left(a_{\cdot,j} - \hat{a}_j(A_{\setminus j})\right)$ is the least squares prediction of the linear predictor as a function of the residuals of $\hat{a}_j$ and $U_j$ is that proportion of the contribution of $a_{\cdot,j}$ to the variance of the linear predictor that cannot be represented as a linear combination of all the other linear terms. We can generalize the unique variance partitioning measure to groups of linear terms as

$$U_l = \frac{\mathbf{1}'_{|B_l|}\mathrm{Var}\left[\hat{\eta}\left(A_{B_l} - \hat{A}_{B_l}(A_{\setminus B_l})\right)\right]\mathbf{1}_{|B_l|}}{\mathrm{Var}[\eta]}$$

where $A_{B_l}$ is matrix with linear terms of the $l$th group as its columns, while the matrix $A_{\setminus B_l}$ collects those terms not in the $l$th group. $\hat{A}_{B_l}(A_{\setminus B_l})$ is the multivariate least-squares predictor for the linear term in the $l$th group. This generalizes the squared semi-partial correlation coefficient to groups of linear terms so that shared variation within the group that cannot be predicted from linear terms in the other groups is incorporated into the unique variance partitioning measure.

The unique variance partition is simple to calculate as it can be written as a function of the covariances between the linear terms in $A$ using the partial variance of $A_{B_l}$ conditional on $A_{\setminus B_l}$ (Whittaker 1990 pp. 134–137, *cf*. Mulaik 2010, pp. 95–100)

$$U_l = \frac{\mathbf{1}'_{|B_l|}\left(\mathrm{Var}[A_{B_l}] - \mathrm{Cov}[A_{B_l}, A_{\setminus B_l}]\mathrm{Var}[A_{\setminus B_l}]^{-1}\mathrm{Cov}[A_{\setminus B_l}, A_{B_l}]\right)\mathbf{1}_{|B_l|}}{\mathrm{Var}[\eta]}$$

$$U_l = \frac{\mathbf{1}'_{|B_l|}\left(K_{A_{B_l}} - K_{A_{B_l},A_{\setminus B_l}}K^{-1}_{A_{\setminus B_l}}K_{A_{\setminus B_l},A_{B_l}}\right)\mathbf{1}_{|B_l|}}{\mathrm{Var}[\eta]}$$

The same approach can be extended to calculate the unique covariances between linear terms or groups of linear terms, but we do not consider them here further (see Whittaker 1990, pp. 134–144).

We outline the proof of the above equations next. From the properties of the least squares predictor, the bilinearity of covariance and the definition of the linear predictor as $\eta = A\mathbf{1}_d$ we get the equality $\mathrm{Var}\left[\hat{\eta}\left(a_{\cdot,j} - \hat{a}_j(A_{\setminus j})\right)\right] = \mathrm{Var}[a_{\cdot,j} - \hat{a}_j(A_{\setminus j})]$. Using notation for partial variance and covariance (Whittaker 1990, pp. 134-138) we can write the two variance terms as

$$\mathrm{Var}[a_{\cdot,j} - \hat{a}_j(A_{\setminus j})] = \mathrm{Var}[a_{\cdot,j}|A_{\setminus j}] = \mathrm{Var}[a_{\cdot,j}] - \mathrm{Cov}[a_{\cdot,j}, A_{\setminus j}]\mathrm{Var}[A_{\setminus j}]^{-1}\mathrm{Cov}[A_{\setminus j}, a_{\cdot,j}]$$

$$\mathrm{Var}\left[\hat{\eta}\left(a_{\cdot,j} - \hat{a}_j(A_{\setminus j})\right)\right] = \mathrm{Cov}[\eta, a_{\cdot,j}|A_{\setminus j}]\mathrm{Var}[a_{\cdot,j}|A_{\setminus j}]^{-1}\mathrm{Cov}[a_{\cdot,j}\eta|A_{\setminus j}]$$

To show that they are equal, we must show that $\mathrm{Cov}[\eta, a_{\cdot,j}|A_{\setminus j}] = \mathrm{Var}[a_{\cdot,j}|A_{\setminus j}]$ in which case

$$\mathrm{Var}\left[\hat{\eta}\left(a_{\cdot,j} - \hat{a}_j(A_{\setminus j})\right)\right] = \mathrm{Var}[a_{\cdot,j}|A_{\setminus j}]\mathrm{Var}[a_{\cdot,j}|A_{\setminus j}]^{-1}\mathrm{Var}[a_{\cdot,j}|A_{\setminus j}] = \mathrm{Var}[a_{\cdot,j}|A_{\setminus j}]$$

The partial covariance is defined as $\mathrm{Cov}[a_{\cdot,j}\eta|A_{\setminus j}] = \mathrm{Cov}[\eta, A_{\setminus j}] - \mathrm{Cov}[\eta, A_{\setminus j}]\mathrm{Var}[A_{\setminus j}]^{-1}\mathrm{Cov}[A_{\setminus j}, a_{\cdot,j}]$.

To simplify presentation, we treat the two additive terms separately and utilize the bilinearity of covariance (Whittaker 1990, pp. 120–124):

$$\begin{aligned}
\mathrm{Cov}[\eta, A_{\setminus j}] &= \mathrm{Cov}[A\mathbf{1}_d, a_{\cdot,j}] \\
\mathrm{Cov}[\eta, A_{\setminus j}] &= \mathrm{Cov}[a_{\cdot,j} + A_{\setminus j}\mathbf{1}_{d-1}, a_{\cdot,j}] \\
\mathrm{Cov}[\eta, A_{\setminus j}] &= \mathrm{Var}[a_{\cdot,j}] + \mathbf{1}'_{d-1}\mathrm{Cov}[A_{\setminus j}, a_{\cdot,j}]
\end{aligned}$$

$$\begin{aligned}
\mathrm{Cov}[\eta, A_{\setminus j}]\mathrm{Var}[A_{\setminus j}]^{-1}\mathrm{Cov}[A_{\setminus j}, a_{\cdot,j}] &= \mathrm{Cov}[a_{\cdot,j} + A_{\setminus j}\mathbf{1}_{d-1}, A_{\setminus j}]\mathrm{Var}[A_{\setminus j}]^{-1}\mathrm{Cov}[A_{\setminus j}, a_{\cdot,j}] \\
\mathrm{Cov}[\eta, A_{\setminus j}]\mathrm{Var}[A_{\setminus j}]^{-1}\mathrm{Cov}[A_{\setminus j}, a_{\cdot,j}] &= \mathrm{Cov}[a_{\cdot,j} + A_{\setminus j}\mathbf{1}_{d-1}, A_{\setminus j}]\mathrm{Var}[A_{\setminus j}]^{-1}\mathrm{Cov}[A_{\setminus j}, a_{\cdot,j}] \\
\mathrm{Cov}[\eta, A_{\setminus j}]\mathrm{Var}[A_{\setminus j}]^{-1}\mathrm{Cov}[A_{\setminus j}, a_{\cdot,j}] & \\
= \mathrm{Cov}[a_{\cdot,j}, A_{\setminus j}]\mathrm{Var}[A_{\setminus j}]^{-1}\mathrm{Cov}[A_{\setminus j}, a_{\cdot,j}] &+ \mathrm{Cov}[A_{\setminus j}\mathbf{1}_{d-1}, A_{\setminus j}]\mathrm{Var}[A_{\setminus j}]^{-1}\mathrm{Cov}[A_{\setminus j}, a_{\cdot,j}]
\end{aligned}$$

Now we only need to combine and reorder the terms:

$$\begin{aligned}
\mathrm{Cov}[a_{\cdot,j}\eta|A_{\setminus j}] =& \ \mathrm{Var}[a_{\cdot,j}] - \mathrm{Cov}[a_{\cdot,j}, A_{\setminus j}]\mathrm{Var}[A_{\setminus j}]^{-1}\mathrm{Cov}[A_{\setminus j}, a_{\cdot,j}] + \mathbf{1}'_{d-1}\mathrm{Cov}[A_{\setminus j}, a_{\cdot,j}] \\
& - \mathrm{Cov}[A_{\setminus j}\mathbf{1}_{d-1}, A_{\setminus j}]\mathrm{Var}[A_{\setminus j}]^{-1}\mathrm{Cov}[A_{\setminus j}, a_{\cdot,j}] \\
\mathrm{Cov}[a_{\cdot,j}\eta|A_{\setminus j}] =& \ \mathrm{Var}[a_{\cdot,j}|A_{\setminus j}] + \mathbf{1}'_{d-1}\mathrm{Cov}[A_{\setminus j}, a_{\cdot,j}] - \mathrm{Cov}[A_{\setminus j}\mathbf{1}_{d-1}, A_{\setminus j}]\mathrm{Var}[A_{\setminus j}]^{-1}\mathrm{Cov}[A_{\setminus j}, a_{\cdot,j}] \\
\mathrm{Cov}[a_{\cdot,j}\eta|A_{\setminus j}] =& \ \mathrm{Var}[a_{\cdot,j}|A_{\setminus j}] + \mathbf{1}'_{d-1}\mathrm{Cov}[A_{\setminus j}, a_{\cdot,j}] - \mathbf{1}'_{d-1}\mathrm{Var}[A_{\setminus j}]\mathrm{Var}[A_{\setminus j}]^{-1}\mathrm{Cov}[A_{\setminus j}, a_{\cdot,j}]
\end{aligned}$$

The latter terms cancel out concluding our proof of $\mathrm{Cov}[\eta, a_{\cdot,j}|A_{\setminus j}] = \mathrm{Var}[a_{\cdot,j}|A_{\setminus j}]$.

#### 2.6.2 Marginal variance partition

Here the term *marginal variance partition* $M_x$ is a measure of variance partitioning with a long history and multiple names (see e.g., Chase 1960, Pratt 1987, Bring 1995). The "marginal" in the name refers both to it being a measure of the direct and the overall effect of a linear term on the variability of the linear predictor (Nathans *et al.* 2012) and to the fact that it can be calculated as either the row or column sum of $K_A$:

$$M_j = \frac{\sum_{j'=1}^{d} K_{A\,j,j'}}{\mathrm{Var}[\eta]}$$
$$M_l = \frac{\sum_{l'=1}^{m} K_{AB\,l,l'}}{\mathrm{Var}[\eta]}$$

The unscaled measure $\sum_{j'=1}^{d} K_{A\,j,j'}$ is equal to the covariance between the linear term and the linear predictor as above. Using again methods from linear regression for our proof, remember that the regression weights can be estimated from the covariance matrix of the covariates and the covariance between the covariates and the outcome variable, that is $\beta' = \mathrm{Cov}[X, Y]'\mathrm{Var}[X]^{-1}$ and hence $\mathrm{Var[X]}\beta = \mathrm{Cov}[X, Y]$ (Whittaker 1990, pp. 124–128, Mulaik 2010, pp. 93 – 95). In our case this corresponds to $\mathrm{Cov}[\eta, A] = K_A\mathbf{1}_d$ as the regression weights are all 1 by definition.

### 2.7 Bayesian inference

The linear predictor and the linear terms are functions of data $D$ and model parameter values $\theta$. In the Bayesian context this implies that they are random variables themselves and that their distribution is a function of the distribution of the parameters, $f_\Theta(\theta)$. However, the distribution of a function of a random variable, say the distribution of the matrix of linear terms, $f_A(\theta)$, is often difficult to express even when the distribution of $\Theta$ is known (Casella and Berger 2002, pp. 47–55). Fortunately, expectations of such functions are well defined in terms of the distribution of $\Theta$. Let $A = g(\theta, D)$, then the posterior expectation of $\mathrm{E}[A|\Theta]$ is $\mathrm{E}[g(\Theta, D)] = \int g(\theta, D)\, f_\Theta(\theta|D) d\theta$ (Casella and Berger 2002, pp. 55–59). This carries over to practical applications using Markov Chain Monte Carlo for Bayesian inference. For samples $\theta_q$ from the posterior distribution of $\Theta$ the mean of function $g(\theta, D)$, $\frac{1}{Q}\sum_{q=1}^{Q} g(\theta_q, D)$ approaches $\mathrm{E}[g(\Theta, D]$ as $Q \to \infty$ for any "well-behaved" Markov chain (Casella and Berger 2002, pp. 269–270). This enables the calculation expectations of any functions of the parameters, such and means and variances of the variance partitions, using the posterior samples.

### 2.8 Variance partitioning in linear models

In the case of linear models, the linear predictor can be also written as a weighted combination of the covariates, $\eta_i = \sum_{j=1}^{d} x_{i,j}\beta_j$ or equivalently $\eta = A\mathbf{1} = X\beta$. Here $x_{i,j} = X_{i,j}$ is the $i$th observations for the $j$th covariate, $x_{\cdot,j}$, and $\beta_j$ is the corresponding regression coefficient. Now $\mathrm{Var}[\eta] = \mathrm{Var}[X\beta] = \beta'\mathrm{Var}[X]\beta$.
