## Supplementary material for "Doors and corners of variance partitioning in statistical ecology": Programming supplement

Variance partitioning tutorial

**Foreword** To simplify the presentation of the essential material, some of the code involved in generating this tutorial has been omitted from the rendered version. Specifically, the accompanying RMarkdown source file can be consulted to see the code for generating the tables and figures. In general, the code in this tutorial aims to demonstrate that the computations involved in working with posterior distributions of variance partitions are rather simple. The code is not written with performance, numerical stability, or customizability in mind, but just to help the applied ecologist understand the kinds of computational tasks involved in variance partitioning.

### Example model: Linearized Ricker model

We demonstrate the use of the variance partitioning methods with a worked example of a simple linear model.

For the demonstration we use the Bayesian regression modeling R-package `rstanarm`. Though less versatile than the Bayesian modelling package `brms`, its syntax is very similar to traditional R regression packages and libraries.

#### R dependencies

```
library('rstanarm')
library('Matrix')
```

#### Data

We use for the demonstration the same Glanville fritillary dataset as in the main study. The corresponding file in the datadryad dataset is `data/survey_data.tsv` inside the archive `ECOG-04799.zip` (Schulz *et al.* 2019, 2020).

```
dat <- read.delim('survey_data.tsv', na.strings = 'NULL')
```

For the demonstration we select a sample of twenty patches that have been continuously inhabited by the butterfly from 1998 to 2009. We split the patches into two groups of ten patches, one for fitting the model and the other for predictive variance partitioning.

```
sample_patches <- c(
       4,    6,   29,   41,  278,  503,  511,  875,  954,  975
)
prediction_patches <- c(
    1051, 1071, 1183, 1189, 1377, 1387, 1642, 1674, 9563, 9668
)

dat <- dat[dat$year < 2010,]
dat$vegetation <- pmax(dat$plantago, dat$veronica)
```

```r
dat_pred <- dat[dat$patch %in% prediction_patches,]
dat <- dat[dat$patch %in% sample_patches,]
```

##### Regression model and inference

We model the (log) growth rate using the linearized Ricker model (see, e.g., Weigell et al., 2021) and fit it using `rstanarm`:

$$\overbrace{\log(\frac{N_{i,t}}{N_{i,t-1}})}^{\text{growth rate}} = y_{i,t} = \overbrace{\alpha + \sum_{j=1}^{d} \beta_j x_{i,j} + \gamma_t}^{\text{intrinsic growth rate}} - \overbrace{\beta_N N_{i,t-1}}^{\text{density dependence}} + \overbrace{\epsilon_{i,t}}^{\text{residual}} .$$

$\gamma_t$ in the *intrinsic growth rate* component of the model corresponds to a yearly random effect and the $x$s to the covariates. Here, we include only host plant abundance and grazing intensity as habitat quality covariates for predicting the growth rate.

```r
dat$growth_rate <- dat$population / dat$previous_population

fit  <- stan_lmer(
    log(growth_rate) ~
        log1p(vegetation) + grazing_presence + (1|year) + previous_population,
    dat
)
```

#### Variance partitioning in practice

In practical applications the most challenging part of implementing variance partitioning as part of the analysis is the forming of the matrix of linear terms and calculation of the covariance matrix for the linear terms. The challenges stem from the need to tailor this step to whatever model structure and software is in use. The general steps involve combining the model covariates and parameters to form a matrix where each column corresponds to one linear term and each row to one observation. For linear regression models constructing the matrix of linear terms amounts to multiplying the covariates by their corresponding regression weights, but models with non-linear and random effects require more effort.

##### Matrix of linear terms

To form the matrix of linear terms, $A$ (See the Mathematical supplement), when working with a (generalized) linear model we can use the design matrix, $X$, of the model. And in this case where the model response is linear we can also use the dependent variable $y$ to calculate the residuals. It can be helpful to avoid forming and storing the matrix of linear terms explicitly, as this will require memory in proportion to the product of the number of observations, number of linear terms and number of posterior samples (*iterations* in the code). Instead it can be sensible to work with the covariance matrix directly when calculating the variance partitioning measures.

The following steps depend on the specific packages used to fit the model, and must be adapted when doing inference with a different tool.

```r
X <- fit$x
X <- X[,!grepl('_NEW', colnames(X))]
y <- fit$y
```

We need the parameter samples, $\theta$ from the posterior distribution as well. For objects of class `stanfit`, this is rather simple:

```
params <- as.matrix(fit)
```

For calculating the different variance partitioning measures for groups of linear terms we need to define the grouping (see below and the Mathematical supplement).

```
groups <- list(
    habitat_quality = c('log1p(vegetation)', 'grazing_presence'),
    density_dependence = c('previous_population'),
    year = 'year',
    residuals = 'residuals'
)

linear_terms <- unname(unlist(groups))
```

To simplify the code and improve readability, we can define auxliary constants for the number of linear terms, number groups of linear terms, and number of posterior samples (or *iterations*).

```
n_lt <- length(linear_terms)
n_group <- length(groups)
n_iter <- nrow(params)
```

We get the linear terms corresponding to the model covariates by simply multiplying the corresponding columns in the design matrix by a diagonal matrix containing the regression weights along the diagonal.

In this case the levels of the *iid* random effect are represented using dummy coding. Hence, to form the linear term for the the yearly *iid* random effect $\gamma_t$, we can collapse the corresponding columns in the design matrix by multiplying with the respective posterior samples for the levels of the random effect.

```
construct_lin_term_mat <- function(params, X, y) {

    ## Linear terms for the growth rate and density dependence
    X_cov <- X[, !grepl('(Intercept)', colnames(X), fixed = TRUE)]
    A_cov <- as.matrix(X_cov %*% Diagonal(x = params[colnames(X_cov)]))
    colnames(A_cov) <- colnames(X_cov)

    ## Linear term for the yearly random effect
    X_iid <- X[,grep('b[(Intercept)', colnames(X), fixed = TRUE)]
    A_iid <- X_iid %*% params[colnames(X_iid)]
    colnames(A_iid) <- c('year')

    A <- cbind(A_cov, A_iid)

    ## Linear term for residuals (omiting the intercept above does not affect variance)
    A <- cbind(A, residuals = y - rowSums(A))

    return(as.matrix(A))
}
```

#### Covariance matrix

The sample covariance matrix of the linear terms, $K_A$, can be used to calculate all the variance partitioning measures presented in this paper (and many others; see the Mathematical supplement). Hence, it is the only thing we need to store for further computations (with the exception of conditional variance partitioning; see below). To avoid storing the matrix of linear terms, it is constructed inside a function that calculates the covariance matrix.

```
lin_term_cov <- function(params, X, y) {
    A <- construct_lin_term_mat(params, X, y)
    A <- A[,linear_terms]
    K <- cov(A)
    dimnames(K) <- list(colnames(A), colnames(A))
    return(K)
}
```

Now we can calculate the posterior distribution of the sample covariance matrix of the linear terms by applying the above function to each sample of posterior parameters. As the `apply` function collapses the output of the function over which it is called to a single dimension, we have to restore the proper dimensions.

```
K <- apply(params, 'iterations', lin_term_cov, X, y)

dim(K) <- c(n_lt, n_lt, n_iter)
dimnames(K) <- list(rows = linear_terms, cols = linear_terms, iterations = NULL)
```

###### Variance partition $V_x$

**Normalizing term** Here, we normalize the variance partitions using the variance of the dependent variable $y$ which is fully observed and, hence, the variance partitioning becomes $V_j = \frac{K_{A_{j,j}}}{\mathrm{Var}[y]}$.

```
V_x <- apply(K, 'iterations', diag) / var(y)
dimnames(V_x) <- list(variance = linear_terms, iterations = NULL)
```

Note though, that for non-linear models the normalizing term is the variance of the linear predictor $\eta$ (see Section 2.2 in the article). The linear predictor $\eta$ would differ for each posterior sample so that the division by $\mathrm{Var}[\eta] = 1'K1$ (or `var(eta) = sum(K)`) would have had to happen inside the `apply` -call or use elementwise division with the denominator differing for each iteration; for example the $V_x$ above would be `V_x <- apply(K, 'iterations', diag) / apply(K, 'iterations', sum)`. This is the case also when we calculate the diagonal variance partitioning $P_x$ as illustrated below.

**Posterior summaries** To calculate posterior summaries, such as means and variances of the variance partitions we can use the posterior samples; expectations of distributions can be approximated by averages over samples from that distribution. For example, the variance of the yearly random effect, $\mathrm{Var}[V_{x_\gamma}]$, can be calculated as simply as `var(V_x['year',])`.

Table 1: Summary of variance partition measures of the linear terms. $\mathrm{Q}_p[V_x] = \Pr[V_x \leq x] \leq p$ is the posterior quantile of the variance partitioning measure.

| | $\mathrm{E}[V_x]$ | $\mathrm{Var}[V_x]$ | $\mathrm{Q}_{.025}[V_x]$ | $\mathrm{Q}_{.05}[V_x]$ | $\mathrm{Q}_{.5}[V_x]$ | $\mathrm{Q}_{.95}[V_x]$ | $\mathrm{Q}_{.975}[V_x]$ |
|---|---|---|---|---|---|---|---|
| vegetation | 0.075 | 0.002 | 0.010 | 0.016 | 0.068 | 0.154 | 0.172 |
| grazing | 0.010 | 0.000 | 0.000 | 0.000 | 0.005 | 0.038 | 0.050 |
| $N_{t-1}$ | 0.202 | 0.005 | 0.079 | 0.093 | 0.196 | 0.337 | 0.367 |
| year | 0.185 | 0.005 | 0.060 | 0.078 | 0.180 | 0.308 | 0.341 |
| residuals | 0.625 | 0.001 | 0.575 | 0.581 | 0.620 | 0.686 | 0.704 |

##### Variance partitioning for groups of linear terms

###### Variance partition $V_b$

Utilizing the properties of variance of linear combinations, we can calculate the covariance matrix for the groups of linear terms as a simple matrix product. For that we need a matrix $B$ with one column for each group and a row for each linear term. The elements of the matrix are one if the linear term of the corresponding row is in the group of the respective column and zero otherwise (see Mathematical supplement). The R code to form the $B$ matrix utilizes the list of group memberships defined above.

```
B <- sapply(groups, function(x) { as.integer(linear_terms %in% x) })

dimnames(B) <- list(linear_terms = linear_terms, groups = colnames(B))
```

Table 2: The $B$-matrix showing the group memberships of the linear terms.

| | habitat quality | density dependence | year | residuals |
|---|---|---|---|---|
| vegetation | 1 | 0 | 0 | 0 |
| grazing | 1 | 0 | 0 | 0 |
| $N_{t-1}$ | 0 | 1 | 0 | 0 |
| year | 0 | 0 | 1 | 0 |
| residuals | 0 | 0 | 0 | 1 |

Given $B$ we can calculate the covariance matrix at the level of groups of linear terms $K_B = B'K_AB$. Then the calculation of $V_b$ follows analogously to $V_x$ above.

```
K_B <- apply(K, 'iterations', function(x, B) { t(B) %*% x %*% B }, B)

dim(K_B) <- c(n_group, n_group, n_iter)
dimnames(K_B) <- list(rows = names(groups), cols = names(groups), iterations = NULL)

V_b <- apply(K_B, 'iterations', diag) / var(y)
dimnames(V_b) <- list(variance = names(groups), iterations = NULL)
```

Table 3: Summaries of the posterior distribution of the variance partition over groups of linear terms.

| | $E[V_b]$ | $Var[V_b]$ | $Q_{.025}[V_b]$ | $Q_{.05}[V_b]$ | $Q_{.5}[V_b]$ | $Q_{.95}[V_b]$ | $Q_{.975}[V_b]$ |
|---|---|---|---|---|---|---|---|
| habitat quality | 0.085 | 0.002 | 0.017 | 0.024 | 0.079 | 0.166 | 0.188 |
| density dependence | 0.202 | 0.005 | 0.079 | 0.093 | 0.196 | 0.337 | 0.367 |
| year | 0.185 | 0.005 | 0.060 | 0.078 | 0.180 | 0.308 | 0.341 |
| residuals | 0.625 | 0.001 | 0.575 | 0.581 | 0.620 | 0.686 | 0.704 |

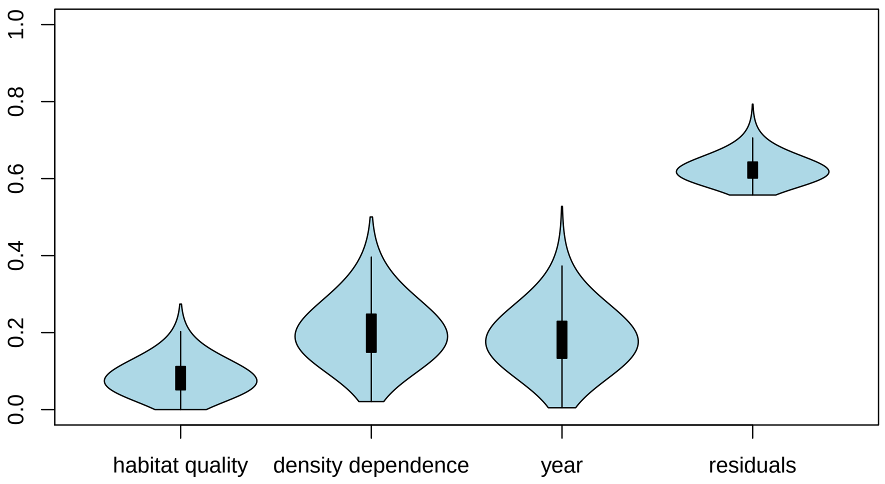

##### Diagonal variance partition $P_b$

In matrix notation the diagonal variance partition is calculated as $V_l = \frac{K_{B_{l,l}}}{\mathrm{tr}(K_B)}$ where tr() is the trace of the matrix, that is the sum of its diagonal terms.

```
P_b <- apply(K_B, 'iterations', function(x) {diag(x) / sum(diag(x))})
dimnames(P_b) <- list(variance = names(groups), iterations = NULL)
```

Table 4: Summaries of the posterior distribution of the diagonal variance partition over groups of linear terms.

| | $\mathrm{E}[P_b]$ | $\mathrm{Var}[P_b]$ | $\mathrm{Q}_{.025}[P_b]$ | $\mathrm{Q}_{.05}[P_b]$ | $\mathrm{Q}_{.5}[P_b]$ | $\mathrm{Q}_{.95}[P_b]$ | $\mathrm{Q}_{.975}[P_b]$ |
|---|---|---|---|---|---|---|---|
| habitat quality | 0.076 | 0.001 | 0.017 | 0.023 | 0.072 | 0.140 | 0.152 |
| density dependence | 0.182 | 0.003 | 0.079 | 0.093 | 0.181 | 0.272 | 0.290 |
| year | 0.167 | 0.003 | 0.060 | 0.075 | 0.165 | 0.262 | 0.279 |
| residuals | 0.576 | 0.004 | 0.459 | 0.477 | 0.570 | 0.694 | 0.718 |

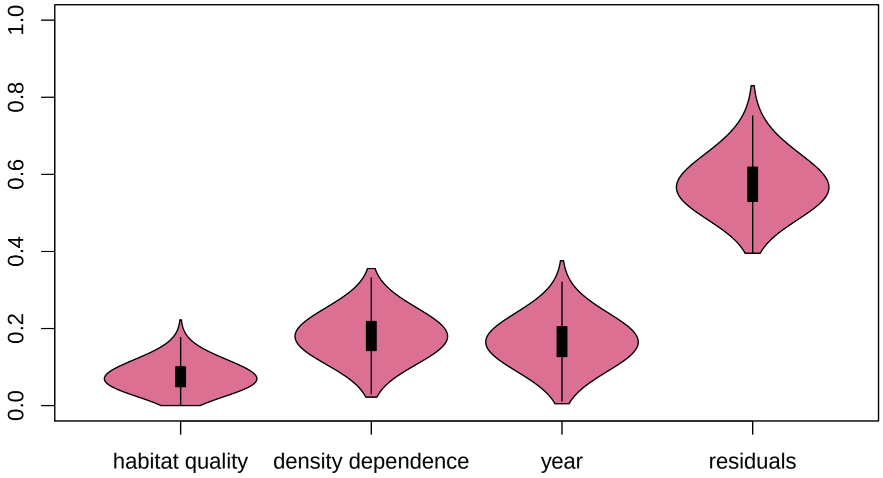


##### Marginal variance partition $M_b$

The marginal variance partition can be formed by taking the row (or column) sums of the covariance matrix.

$$M_l = \frac{(K_B 1_m)_l}{\mathrm{Var}[y]}$$

```
M_b <- apply(K_B, 'iterations', rowSums) / var(y)
dimnames(M_b) <- list(variance = names(groups), iterations = NULL)
```

Table 5: Summaries of the posterior distribution of the marginal variance partition over groups of linear terms.

| | $\mathrm{E}[M_b]$ | $\mathrm{Var}[M_b]$ | $\mathrm{Q}_{.025}[M_b]$ | $\mathrm{Q}_{.05}[M_b]$ | $\mathrm{Q}_{.5}[M_b]$ | $\mathrm{Q}_{.95}[M_b]$ | $\mathrm{Q}_{.975}[M_b]$ |
|---|---|---|---|---|---|---|---|
| habitat quality | 0.041 | 0.000 | 0.015 | 0.020 | 0.041 | 0.062 | 0.066 |
| density dependence | 0.175 | 0.001 | 0.112 | 0.121 | 0.175 | 0.230 | 0.240 |
| year | 0.177 | 0.002 | 0.088 | 0.103 | 0.178 | 0.248 | 0.260 |
| residuals | 0.607 | 0.003 | 0.498 | 0.516 | 0.606 | 0.700 | 0.719 |

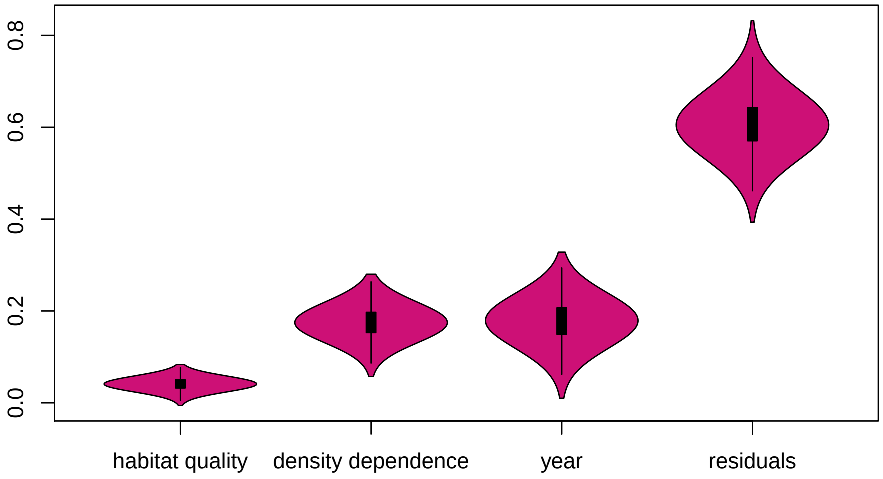


##### Unique variance partition $U_b$

The unique variance partition is calculated as the sum of the residual variance of the least squares prediction of a group of linear terms as a function of the linear terms in all the other groups. In practice this is simple to implement using the sample covariance matrix $K_A$ and the grouping matrix $B$.

$$U_l = \frac{1'_{|B_l|}(K_{A_{B_l,B_l}} - K_{A_{B_l,B_{\setminus l}}} K^{-1}_{A_{B_{\setminus l},B_{\setminus l}}} K_{A_{B_{\setminus l},B_l}})1_{|B_l|}}{\text{Var}[y]}$$

Here $B_l$ is the index of columns in $A$ or $K_A$ that belong to the $l$'th group of linear terms and $B_{\setminus l}$ is its complement, that is the linear terms in all the other groups. $|B_l|$ is the number of linear terms in that group.

The R implementation below uses nested functions, where the innermost calculates $U_b$ for one group of linear terms and the outer function iterates over all linear term groups.

```
unique_variance <- function(K, idx) {
    U <- sum(K[idx,idx] - K[idx,!idx] %*% solve(K[!idx,!idx]) %*% K[!idx,idx])
}

unique_variances <- function(K, B) {
    U <- apply(B, 'groups', function(x, K) { unique_variance(K, x == 1 )}, K)
}

U_b <- apply(K, 'iterations', unique_variances, B) / var(y)
dimnames(U_b) <- list(variance = names(groups), iterations = NULL)
```

Table 6: Summaries of the posterior distribution of the unique variance partition over groups of linear terms.

| | $\text{E}[U_b]$ | $\text{Var}[U_b]$ | $\text{Q}_{.025}[U_b]$ | $\text{Q}_{.05}[U_b]$ | $\text{Q}_{.5}[U_b]$ | $\text{Q}_{.95}[U_b]$ | $\text{Q}_{.975}[U_b]$ |
|---|---|---|---|---|---|---|---|
| habitat quality | 0.080 | 0.002 | 0.016 | 0.022 | 0.075 | 0.154 | 0.172 |
| density dependence | 0.191 | 0.005 | 0.073 | 0.087 | 0.186 | 0.314 | 0.338 |
| year | 0.176 | 0.005 | 0.057 | 0.073 | 0.173 | 0.293 | 0.318 |
| residuals | 0.601 | 0.001 | 0.561 | 0.565 | 0.596 | 0.653 | 0.668 |

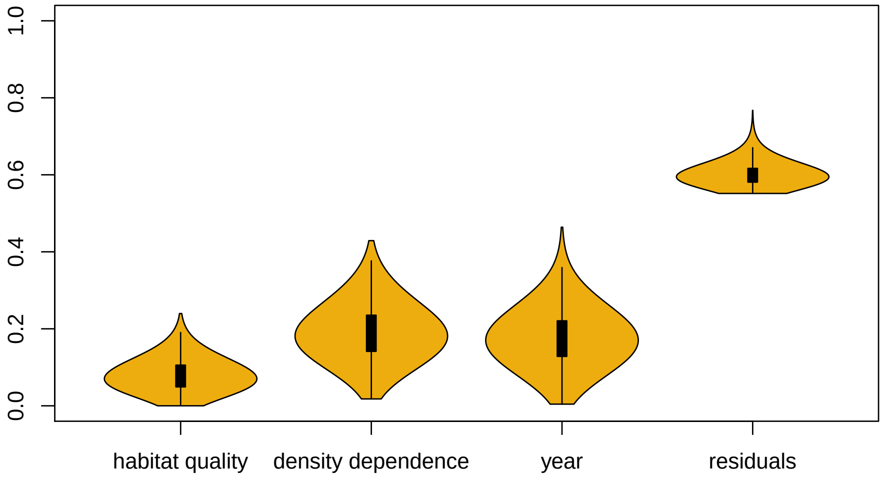


#### Sample, population, and predictive variance partitioning

From a computational perspective, the *sample variance partition* is the variance partition calculated from the data used in model fitting, the *predictive variance partition* is the variance partition calculated for some new data, and the *population variance partition* is the variance partition calculated over both new and old data. Whether or not the "old" and "new" data one uses can meaningfully be associated with a "prediction" or a "population" is scientifically relevant, but has no bearing on implementing the computations.

##### Predictive variance partition

For predictive variance partitioning we need covariates for the "prediction scenario", or in our case the set of ten patches that we excluded from model inference. As we construct the design matrix outside of the modeling framework of `rstanarm` we need to take care and apply all data transformations that were part of the model specification.

```
dat_pred$vegetation <- log1p(dat_pred$vegetation)

y_pred <- log(dat_pred$population / dat_pred$previous_population)
X_pred <- dat_pred[, c('vegetation', 'grazing_presence', 'previous_population')]
colnames(X_pred) <- c('log1p(vegetation)', 'grazing_presence', 'previous_population')
```

Here we implement the dummy coding for the yearly random effect on our own, but generally for non-linear and random effects a more robust and comprehensive method of matching the values of the random effects to the new data is required.

```
years <- seq(min(dat_pred$year), max(dat_pred$year))
X_iid <- t(sapply(dat_pred$year, function(x) { as.integer(years %in% x)}))
colnames(X_iid) <- paste0('b[(Intercept) year:', years, ']')

X_pred <- as.matrix(cbind(X_pred, X_iid))
```

Now that we have constructed the design matrix for the prediction, the following steps are identical to sample variance partitioning. For more complex cases the code creating the matrix of linear terms would have to be updated to take the prediction into account. For example, if the prediction data included years not part of the sample data, we would have to sample values for the new years using the *iid* random effect's distribution; see the Mathematical supplement and the Code supplement for the case study results discussed in the article for an example on spatial random effects.

```
K_pred <- apply(params, 'iterations', lin_term_cov, X_pred, y_pred)
```

```
dim(K_pred) <- c(n_lt, n_lt, n_iter)
dimnames(K_pred) <- list(rows = linear_terms, cols = linear_terms, iterations = NULL)

K_B_pred <- apply(K_pred, 'iterations', function(x, B) { t(B) %*% x %*% B }, B)

dim(K_B_pred) <- c(n_group, n_group, n_iter)
dimnames(K_B_pred) <- list(rows = names(groups), cols = names(groups), iterations = NULL)

V_b_pred <- apply(K_B_pred, 'iterations', diag) / var(y)
dimnames(V_b_pred) <- list(variance = names(groups), iterations = NULL)
```

Table 7: Summaries of the posterior distribution of the predicted variance partition over groups of linear terms.

| | $E[V_b^{pred}]$ | $Var[V_b^{pred}]$ | $Q_{.025}[V_b^{pred}]$ | $Q_{.05}[V_b^{pred}]$ | $Q_{.5}[V_b^{pred}]$ | $Q_{.95}[V_b^{pred}]$ | $Q_{.975}[V_b^{pred}]$ |
|---|---|---|---|---|---|---|---|
| habitat quality | 0.035 | 0.000 | 0.007 | 0.009 | 0.031 | 0.075 | 0.087 |
| density dependence | 0.154 | 0.003 | 0.060 | 0.071 | 0.149 | 0.257 | 0.279 |
| year | 0.185 | 0.005 | 0.060 | 0.078 | 0.180 | 0.308 | 0.341 |
| residuals | 0.534 | 0.002 | 0.455 | 0.465 | 0.528 | 0.621 | 0.643 |

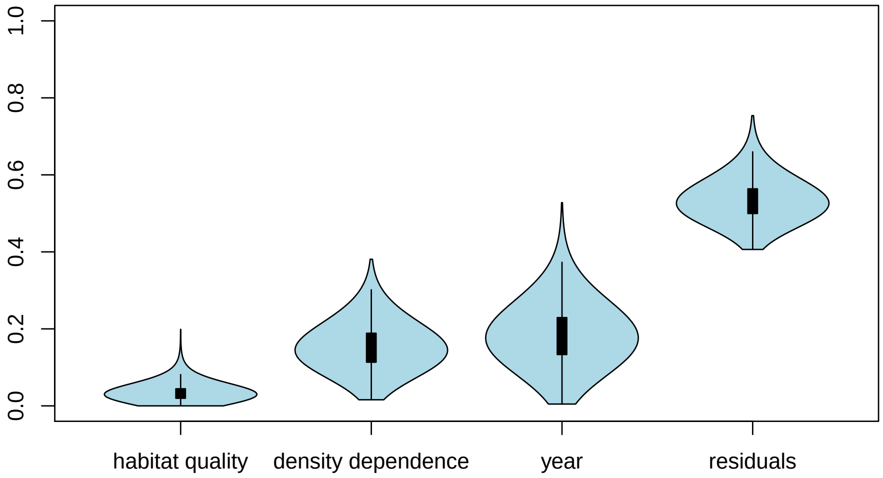


###### Population variance partition

For the population variance partitioning we only need to concatenate the data and the we can proceed as usual.

```
X_pop <- rbind(X[,colnames(X_pred)], X_pred)
y_pop <- c(y, y_pred)

K_pop <- apply(params, 'iterations', lin_term_cov, X_pop, y_pop)

dim(K_pop) <- c(n_lt, n_lt, n_iter)
dimnames(K_pop) <- list(rows = linear_terms, cols = linear_terms, iterations = NULL)

K_B_pop <- apply(K_pop, 'iterations', function(x, B) { t(B) %*% x %*% B }, B)

dim(K_B_pop) <- c(n_group, n_group, n_iter)
dimnames(K_B_pop) <- list(rows = names(groups), cols = names(groups), iterations = NULL)
```

```
V_b_pop <- apply(K_B_pop, 'iterations', diag) / var(y)
dimnames(V_b_pop) <- list(variance = names(groups), iterations = NULL)
```

Table 8: Summaries of the posterior distribution of the population variance partition over groups of linear terms.

| | $E[V_b^{pop}]$ | $Var[V_b^{pop}]$ | $Q_{.025}[V_b^{pop}]$ | $Q_{.05}[V_b^{pop}]$ | $Q_{.5}[V_b^{pop}]$ | $Q_{.95}[V_b^{pop}]$ | $Q_{.975}[V_b^{pop}]$ |
|---|---|---|---|---|---|---|---|
| habitat quality | 0.061 | 0.001 | 0.013 | 0.017 | 0.056 | 0.117 | 0.132 |
| density dependence | 0.177 | 0.004 | 0.070 | 0.082 | 0.172 | 0.296 | 0.322 |
| year | 0.184 | 0.005 | 0.060 | 0.078 | 0.179 | 0.307 | 0.339 |
| residuals | 0.578 | 0.001 | 0.528 | 0.533 | 0.574 | 0.637 | 0.651 |

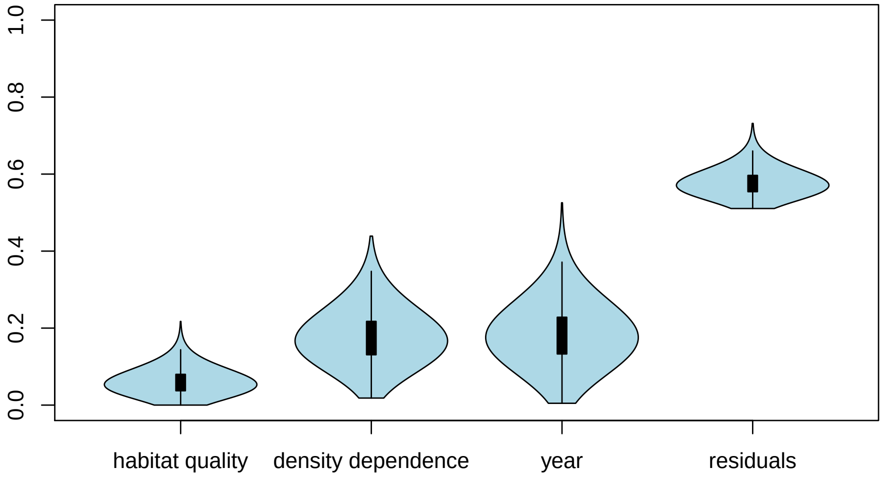


#### Conditional variance partitioning

For conditional variance partitioning we choose as the conditioning variable the abundance of the host plant which in this case is an ordinal variable (`vegetation`).

```
g <- dat$vegetation
G <- unique(g)

n_cond <- length(G)
```

##### Conditional variance partitioning

The conditional variance partition is simply the variance partition calculate separately over the observation falling into each level of the conditioning variable.

```
conditional_lin_term_cov <- function(params, G, g, X, y) {
    A <- construct_lin_term_mat(params, X, y)

    K <- t(sapply(G, function(x, A, g) { cov(A[g == x,])}, A, g))
    dim(K) <- c(length(G), rep(ncol(A), 2))
    dimnames(K) <- list(condition = G, colnames(A), colnames(A))

    return(K)
}
```

The calculations start again from the covariance matrix, now one for each value of `vegetation`. The rest of the code differs only in the multiple covariance matrices per posterior sample, and the normalizing variance term, which is calculate only with respect to observations within the same condition.

```
K_V <- apply(params, 'iterations', conditional_lin_term_cov, G, g, X, y)

dim(K_V) <- c(n_cond, n_lt, n_lt, n_iter)
dimnames(K_V) <- list(
    condition = G,
    rows = linear_terms,
    cols = linear_terms,
    iterations = NULL
)

K_V <- (apply(K_V, c('condition', 'iterations'), function(x, B) { t(B) %*% x %*% B }, B))

dim(K_V) <- c(n_group, n_group, n_cond, n_iter)
dimnames(K_V) <- list(
    rows = names(groups),
    cols = names(groups),
    condition = G,
    iterations = NULL
)

V_cond <- apply(K_V, c('condition', 'iterations'), function(x) {diag(x) / sum(x)})
dimnames(V_cond) <- list(variance = groups_pretty, condition = G, iterations = NULL)
```

Table 9: Summaries of the posterior distribution of the conditional variance partition over groups of linear terms.

| | $E[V_{\text{veg}=3}]$ | $E[V_{\text{veg}=2}]$ | $E[V_{\text{veg}=1}]$ |
|---|---|---|---|
| habitat quality | 0.01 | 0.01 | 0.02 |
| density dependence | 0.24 | 0.02 | 0.01 |
| year | 0.19 | 0.16 | 0.08 |
| residuals | 0.62 | 0.65 | 0.60 |

##### Variance within and between conditions

Using the set of possible values that `vegetation` can have, and the values of the variable for the observations, we create the matrix $H$, which using dummy coding assings each observation to one level of the conditioning variable.

```
H <- sapply(G, function(x, g) { as.integer(g == x) }, g)
colnames(H) <- G
```

The matrix $H$ can be used to generate a matrix of between-condition linear term means, which we use to calculate the covariance matrix for both within and between conditions (see also Mathematical supplement).

```
wit_btw_lin_term_cov <- function(params, H, X, y) {
    A <- construct_lin_term_mat(params, X, y)

    A_cm <- H %*% diag(1 / colSums(H)) %*% t(H) %*% A

    K_wit <- cov(A - A_cm)
```

```
    K_btw <- cov(A_cm)

    K <- abind::abind(K_wit, K_btw, rev.along = 0)
    dimnames(K) <- list(
        rows = colnames(A),
        cols = colnames(A),
        partition = c('within', 'between')
    )

    return(K)
}
```

Again, the calculation of the variance partition follows a familiar pattern, only there are two covariance matrices for each posterior sample. Here the normalizing variance is again the total variance over all observation.

```
K_WB <- apply(params, 'iterations', wit_btw_lin_term_cov, H, X, y)

dim(K_WB) <- c(n_lt, n_lt, 2, n_iter)
dimnames(K_WB) <- list(
    rows = linear_terms,
    cols = linear_terms,
    partition = c('within', 'between'),
    iterations = NULL
)

K_WB <- apply(K_WB, c('partition', 'iterations'), function(x, B) { t(B) %*% x %*% B }, B)

dim(K_WB) <- c(n_group, n_group, 2, n_iter)
dimnames(K_WB) <- list(
    rows = names(groups),
    cols = names(groups),
    partition = c('within', 'between'),
    iterations = NULL
)

V_wb <- apply(K_WB, c('partition', 'iterations'), function(x) {diag(x)}) /  var(y)
dimnames(V_wb) <- list(
    variance = groups_pretty,
    partition = c('within', 'between'),
    iterations = NULL
)
```

Table 10: Summaries of the posterior distribution of the variance partition over groups of linear terms within and between conditions.

| | $E[V^{\text{wit}}]$ | $E[V^{\text{btw}}]$ |
|---|---|---|
| habitat quality | 0.01 | 0.07 |
| density dependence | 0.19 | 0.01 |
| year | 0.18 | 0.00 |
| residuals | 0.61 | 0.01 |

#### Correlations and covariances

##### Posterior Correlations between variance partition measures

To look at posterior correlations (or covariances) between the variance partitions, one simply calculates the correlation between the posterior sample from the variance partitions of the different groups of linear terms.

```
cor_V_b  <- cor(aperm(V_b, c('iterations', 'variance')))
dimnames(cor_V_b) <- list(rows = groups_pretty, cols = groups_pretty)
```

Table 11: Posterior correlations between the variance partitions.

| | habitat quality | density dependence | year | residuals |
|---|---|---|---|---|
| habitat quality | 1.00 | 0.15 | 0.08 | 0.03 |
| density dependence | 0.15 | 1.00 | 0.00 | 0.01 |
| year | 0.08 | 0.00 | 1.00 | -0.35 |
| residuals | 0.03 | 0.01 | -0.35 | 1.00 |

##### Covariances between linear terms

To summarize the posterior distribution of covariances (or correlations) between the linear terms one can use the covariance matrices used in the calculation of the variance partitions.

```
cov_B <- apply(K_B, c('rows', 'cols'), mean) / var(y)
dimnames(cov_B) <- list(rows = groups_pretty, cols = groups_pretty)

cor_B <- apply(K_B, 'iterations', cov2cor)
dim(cor_B) <- c(n_group, n_group, n_iter)
dimnames(cor_B) <- list(rows = groups_pretty, cols = groups_pretty, iterations = NULL)

cor_B <- apply(cor_B, c('rows', 'cols'), mean)
```

Table 12: Posterior mean of normalized covariances between the linear terms.

| | habitat quality | density dependence | year | residuals |
|---|---|---|---|---|
| habitat quality | 0.08 | -0.02 | -0.01 | -0.01 |
| density dependence | -0.02 | 0.20 | 0.00 | -0.01 |
| year | -0.01 | 0.00 | 0.18 | 0.00 |
| residuals | -0.01 | -0.01 | 0.00 | 0.62 |

Table 13: Posterior mean of correlations between the linear terms.

| | habitat quality | density dependence | year | residuals |
|---|---|---|---|---|
| habitat quality | 1.00 | -0.19 | -0.07 | -0.03 |
| density dependence | -0.19 | 1.00 | 0.02 | 0.00 |
| year | -0.07 | 0.02 | 1.00 | 0.01 |
| residuals | -0.03 | 0.00 | 0.01 | 1.00 |
