## Supplementary material for "Doors and corners of variance partitioning in statistical ecology": Model supplement

#### 1 SPATIO-TEMPORAL RANDOM EFFECT

For the spatio-temporal random effect $z$, we used the stochastic partial differential equation approximation to a Gaussian random field with a Matérn covariance function as implemented in R-INLA (Lindgren *et al*. 2011, Lindgren and Rue 2015). The approximation requires a mesh triangulation over the spatial domain. We used the 2D-triangulation algorithm provided by R-INLA, using the centroids of the habitat patches as the starting locations. The cut-off for joining nearby vertices was set to 0.75 km and the boundary of the inner domain was set to 10 km from the mesh points and for the outer domain it was 20 km from the boundary. The triangulation parameters were such that the maximum edge length was 15 km on the inside and 30 km in the outer domain. The minimum angle for triangulation was set to 21 degrees. After the initial mesh construction, the mesh inside the survey areas was refined so that the maximum edge length is 5 km and the minimum angle 26 degrees (see the map below).

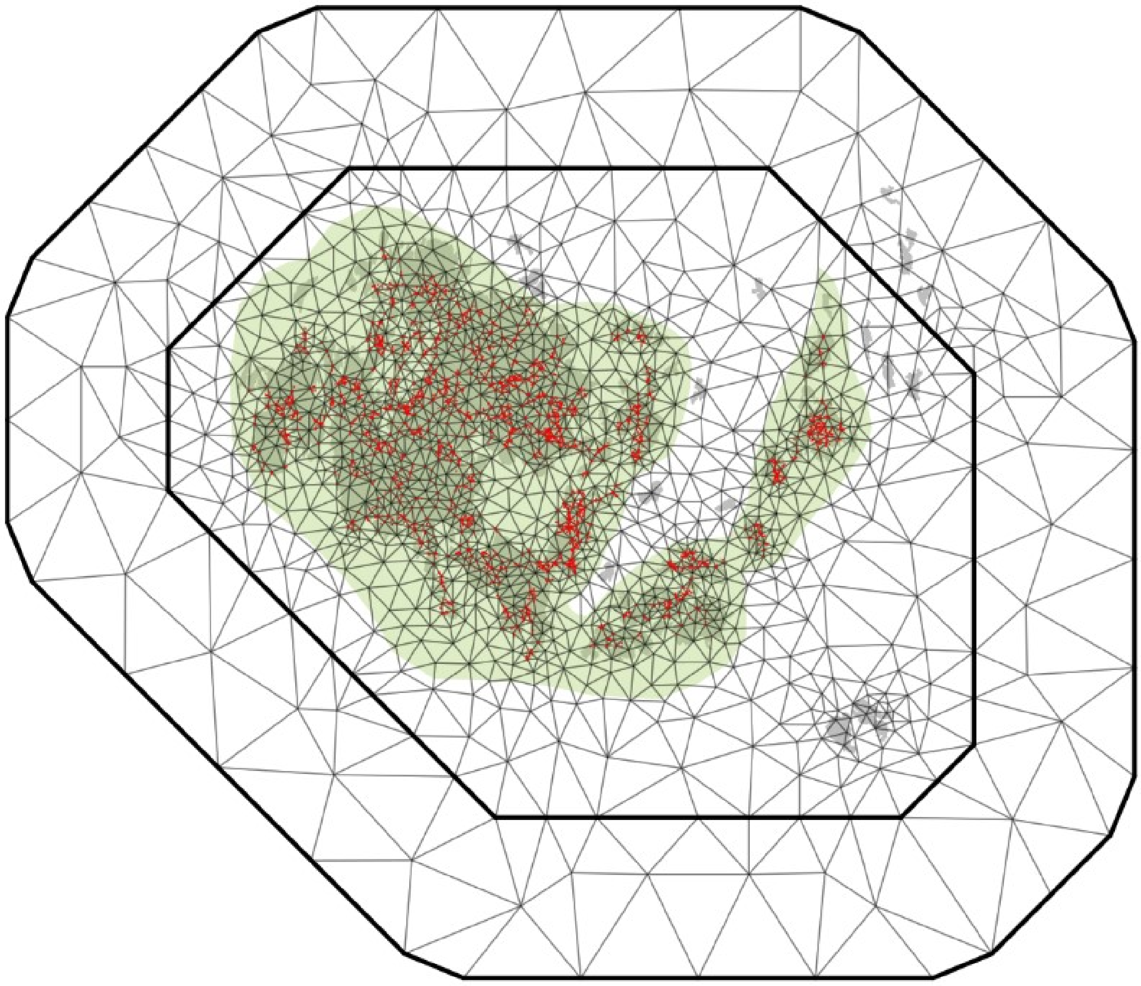

***Figure***. *Map of the triangulation mesh for the SPDE approximation. The green shading indicates the survey areas, which were used in refining the inner mesh domain. The islands in grey use GSHHG data. The number of nodes in the mesh is 1545.*

### 2 PRIOR DISTRIBUTIONS

For the linear covariate effects, we used a Gaussian distribution with mean zero and standard deviation one except for the intercept's prior with standard deviation of 10. The *iid* random effects for patch, $u$, and year, $v$, had priors constructed such that the probability of their standard deviation being over one was 0.5 (see Simpson *et al*. 2017). The range parameter of the spatio-temporal random field had a prior such that the probability of the correlation range being less than one kilometer was 0.05. The variance of the same had a probability of 0.5 for exceeding one (Fuglstad *et al*. 2018). For the temporal autocorrelation parameter in the spatio-temporal random field we used the default prior in R-INLA.

### 3 MODEL COVARIATES

Area, connectivity, and vegetation were log transformed. All covariates were centered around zero and scaled to standard deviation one after the transformation, if any. The covariates and random effects as well as the groups of linear terms they are assigned to are listed below. Additional details about the covariate data are provided in Schulz *et al*. (2020).

***Table***. *Covariate and random effect linear terms in the model and their assignations to groups of linear terms.*

| Covariate / Linear term | Group | Description |
|---|---|---|
| Previous presence | Population | Indicator for occupancy in the previous year |
| Area | Metapopulation | Area of the habitat patch |
| Connectivity | Metapopulation | Expected population connectivity for the habitat patch |
| Vegetation | Habitat quality | Abundance of the larval host plant (ordinal scale) |
| Both hosts | Habitat quality | Indicator variable for the presence of both host plants |
| Dry vegetation | Habitat quality | Proportion of desiccated host plant |
| Grazing intensity | Habitat quality | Proportion of habitat patch grazed |
| Grazing presence | Habitat quality | Indicator of grazing in the habitat patch |
| Patch | Random effect | Per habitat patch *iid* random effect |
| Year | Random effect | Yearly *iid* random effect |
| Spatio-temporal | Random effect | Spatio-temporal random effect |

### 4 POSTERIOR SAMPLING FROM THE JOINT DISTRIBUTION

Posterior samples from the joint distribution of the regression weights and random effects were obtained from R-INLA using the function **inla.posterior.sample**. It generates samples from the joint distribution implied by the Laplace approximation of the latent variables, here the linear covariate effects and the random effects, conditional on model hyperparameters, that is the random effect variances, range and temporal autocorrelation, which are approximated separately by R-INLA (Rue *et al*. 2009). We used 1000 posterior samples for our analyses.

### 5 POSTERIOR PREDICTIVE SAMPLES FOR THE SPATIO-TEMPORAL RANDOM EFFECT

For the spatio-temporal random effect sampling values for the excluded habitat patches' observations is rather straightforward, as our triangulation domain also covered those locations. Conditional on the posterior value of the random field approximation at the mesh nodes, the random effect is simply a linear function of those values (Lindgren and Rue 2015). Hence, we only needed to create a new projection matrix for the unobserved locations and times. Then a posterior sample of the predicted random field values corresponded to the matrix product of projection matrix times the value of the random field approximation in the corresponding posterior sample.

### 6 MARGINAL POSTERIOR DISTRIBUTIONS OF MODEL HYPERPARAMETERS

***Table**. Posterior summaries of the model hyperparameters.*

| **Parameter** | **Mean** | **S.D.** | **95% Cr.I.** |
|---|---|---|---|
| Standard deviation (patch) | 0.79 | 0.06 | 0.69 – 0.92 |
| Standard deviation (year) | 1.10 | 0.19 | 0.79 – 1.53 |
| Standard deviation (space) | 0.52 | 0.08 | 0.36 – 0.69 |
| Correlation distance | 3.37 | 1.61 | 1.43 – 7.54 |
| Autocorrelation | 0.88 | 0.05 | 0.77 – 0.94 |

### 7 MARGINAL POSTERIOR DISTRIBUTIONS OF COVARIATE REGRESSION COEFFICIENTS

***Table**. Posterior summaries of the covariate regression coefficients.*

| **Parameter** | **Mean** | **S.D.** | **95% Cr.I.** |
|---|---|---|---|
| Intercept | -3.43 | 0.26 | -3.94 – -2.92 |
| Previous presence | 0.31 | 0.03 | 0.26 – 0.36 |
| Area | 0.35 | 0.04 | 0.26 – 0.44 |
| Connectivity | 2.02 | 0.09 | 1.84 – 2.19 |
| Both hosts | 0.03 | 0.04 | -0.04 – 0.10 |
| Dry vegetation | 0.04 | 0.03 | -0.02 – 0.10 |
| Grazing intensity | -0.33 | 0.04 | -0.41 – -0.25 |
| Grazing presence | 0.06 | 0.04 | -0.02 – 0.14 |
| Vegetation | 0.94 | 0.04 | 0.86 – 1.01 |
